# Cytochrome P450 inhibition impedes pyrethroid effects on insects through Nav channel regulation

**DOI:** 10.1101/2025.02.03.636193

**Authors:** Beata Niklas, Eléonore Moreau, Caroline Deshayes, Jakub Rydzewski, Milena Jankowska, Jacek Kęsy, Ademir J. Martins, Pie Müller, Vincent Corbel, Wiesław Nowak, Bruno Lapied

**Author notes:** Present address; Univ Angers, BIODIVAG, 49000 Angers, France (Eléonore Moreau). Present address; Univ Angers, Univ Brest, IRF, SFR ICAT, F-49000 Angers, France. Corresponding Author: Professor B. Lapied at the above address. e.mail. These authors contributed equally to this work and share first authorship (Beata Niklas, Eléonore Moreau).

## Abstract

Insecticides used in various formulations are inevitable in agricultural pest control and prevention of vector-borne diseases. Piperonyl butoxide (PBO) is a synergist widely used to enhance the effectiveness of insecticides, notably pyrethroids, by inhibiting the detoxifying cytochrome P450 enzymes, thus reducing the capacity of insects to metabolize and resist insecticides. Recent studies, however, reveal an unexpectedly weak restoration of pyrethroid efficacy by PBO, but the underlying mechanism is unknown. Here, we demonstrate that the PBO-induced inhibition of cytochrome P450 impedes the effect of deltamethrin by mainly affecting its interaction with the inactivated state of voltage-gated sodium channels (Nav). We describe a new octopamine-dependent regulatory mechanism involving Gαs, PKA, DARPP-32, and PP1-2A, which affect the cytochrome P450 conformation, thus limiting the effect of PBO and modulating the Nav gating. As a result, deltamethrin cannot reach its final binding site in the fenestration to exert its full effect. We confirmed *in vivo* that under chemical stressor, the level of octopamine is elevated, which decreases the deltamethrin efficacy. Our findings reveal a novel adaptation mechanism that increases insect survival by reducing insecticide efficacy, thus emphasizing the necessity of developing more effective formulations and technologies for pest and vector control.

## Introduction

Insects, accounting for approximately 80% of the species in the world, possess an impressive capacity to adapt to environmental changes, including biotic, abiotic and physical stressors^1,2^. Although less than 2-3% of insects are pests or harmful, they contaminate food, destroy cultivated crops^3^, and transmit pathogen protozoa and viruses resulting in more than 700,000 deaths each year^4^. The increase in their spread has become significantly more pronounced in recent years due to climate change, increased travel, trade, and intensive farming practices^5–7^. Synthetic or naturally occurring insecticides^8^ are still the cornerstone of pest and vector-borne disease control worldwide^6,9^. Currently, pyrethroids are one of the most important groups of insecticides used to reduce the pest population and are recommended by the World Health Organization (WHO) for protecting humans against mosquito-borne diseases^10,11^. However, the massive use of pyrethroids led to the emergence and spread of pyrethroid resistance through behavioural, physiological, and genetic mechanisms including target site modifications. Insects can develop several of these mechanisms simultaneously^12–14^ hence becoming “super– insecticide-resistant”^15^. Upregulation of detoxifying cytochrome P450 enzymes, responsible for the biotransformation of xenobiotics, is one of the most prevalent mechanisms of pyrethroid resistance^16–18^. To overcome this problem, the cytochrome P450 inhibitor piperonyl butoxide (PBO)^19–21^, has been widely used as a synergist in combination with pyrethroids in classic formulations to enhance their overall effectiveness.

Pyperonyl butoxide, developed in the 1940s, is included in over 2500 insecticide formulations and used in agriculture, veterinary medicine, and the prevention of vector-borne diseases^22–25^. Moreover, PBO has been incorporated recently into pyrethroid-based Long-Lasting Insecticidal Nets (LLIN), which are currently produced by seven LLIN manufacturers^26^ and, following a recommendation from the WHO, are being included in national distribution campaigns.

However, recent reports raise concerns about the actual performance of PBO as a synergist. Previous studies have indicated an unexpectedly weak restoration of pyrethroid efficacy by PBO in susceptible insects^27–31^. In addition, although PBO nets generally perform better than conventionally treated nets in high pyrethroid resistance settings, little evidence is available to support greater entomological efficacy of pyrethroid-PBO nets in areas where mosquitoes show moderate to low levels of resistance to pyrethroids ^24^. These findings question the performance of PBO in enhancing pyrethroid efficacy, which could represent an obstacle to effective, sustainable pest and disease vector control.

To date, no data is available to explain the mechanisms behind this limited performance of PBO. Our study aims to bring new insights into the mode of action of PBO and to examine why its interaction with cytochrome P450 impedes the effect of pyrethroids. Using a multidisciplinary approach, we characterized the signaling pathways involved in the cytochrome P450-mediated regulation of the gating properties of voltage-gated sodium channels (Nav) targeted by pyrethroid insecticides. This regulatory mechanism hampers the efficacy of the gold standard pyrethroid deltamethrin on Nav. To prove the physiological relevance, we showed how octopamine, a biogenic amine produced and released under stress conditions^32^, regulates the cytochrome P450 function, reducing the action of deltamethrin. We thus performed *in vivo* experiments to evaluate the impact of exposure to sublethal doses of insecticide on the octopamine-induced decrease of the deltamethrin effect. These results reveal a novel mechanism of insect adaptation to stress conditions that decreases insecticide efficacy.

## Results

### PBO combined with type II pyrethroid show limited efficacy

We first assessed the sensitivity to type II pyrethroid of two susceptible colonies of mosquito, *Aedes aegypti*, of the Rockefeller and Fiocruz strains (Fig. 1a) using the standard WHO cylinder test procedure ^33^ with few modifications to allow for the assessment of the potentiation effect of the synergist PBO on pyrethroid action. Although PBO (Fig. 1d) increased the toxicity of the type II pyrethroid alpha-cypermethrin at low concentrations (0.1-1%), a plateau of mortality was rapidly reached at higher concentrations (from 1% to 4%) (Fig. 1b, c). Surprisingly, increasing the concentrations of PBO (e.g., 4%, a percentage recommended by WHO in the formulation) reduced the toxicity of alpha-cypermethrin in the two susceptible *Ae. aegypti* colonies.

**Fig. 1.**
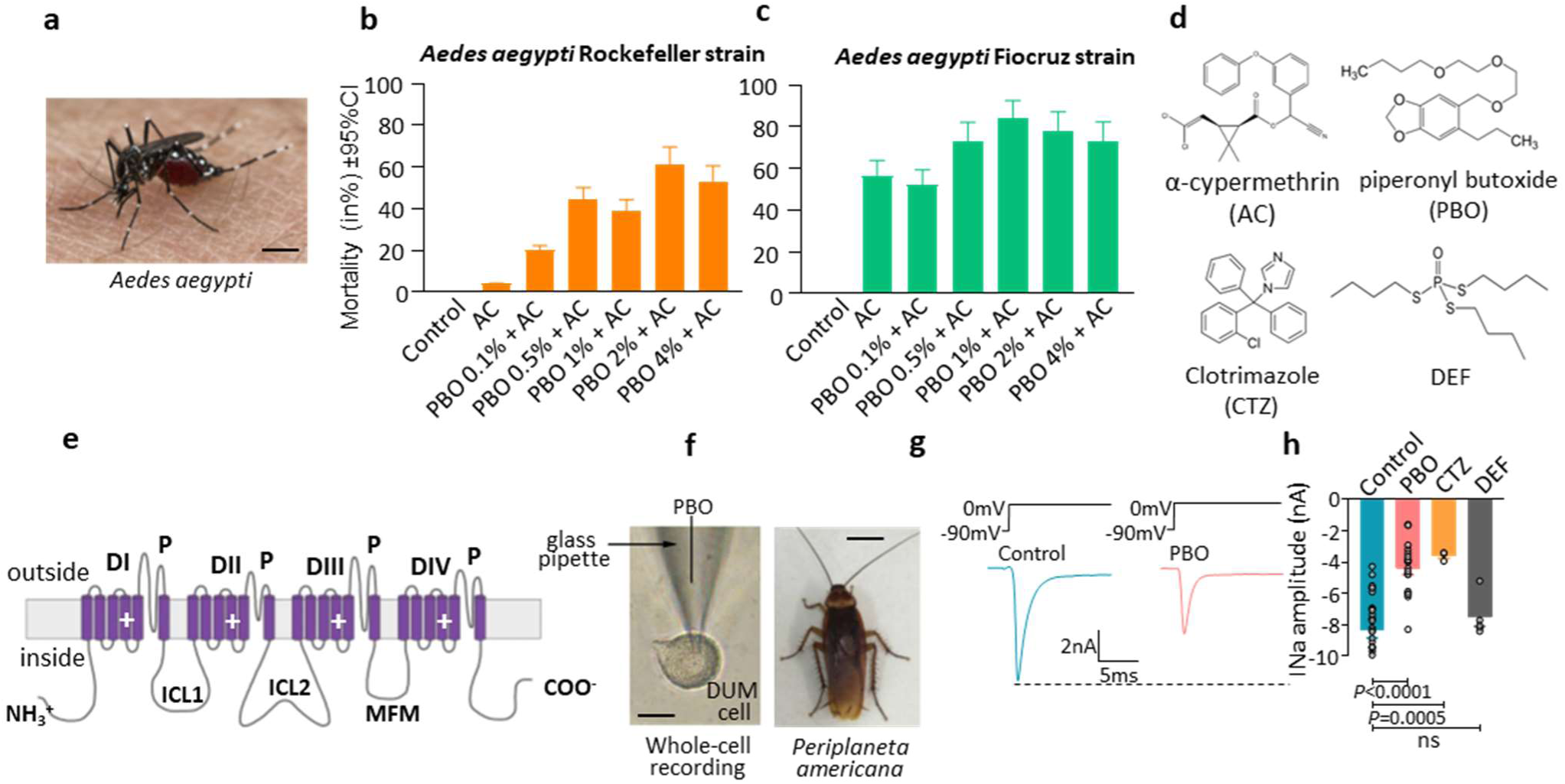
The synergist PBO does not produce full optimized effects of type II pyrethroid against susceptible mosquito strains and alters voltage-gated sodium channels by reducing the inward sodium current amplitude. **a.** Photo of the mosquito *Aedes aegypti*, the main vector of dengue. **b.** Mortality of susceptible *A. aegypti* mosquitoes (Rockefeller strain, Swiss TPH, Switzerland) sequentially exposed for 1h to piperonyl butoxide (PBO, at 0.1–4%) and then for 1h to filter papers impregnated at 0.01% alpha-cypermethrin (type II pyrethroid). Error bars represent 95% confidence intervals. **c.** Mortality of susceptible *A. aegypti* mosquitoes (Rockefeller strain, FIOCRUZ, Brazil) sequentially exposed for 1h to PBO (at 0.1–4%) and then for 1h to filter papers impregnated at 0.02% alpha-cypermethrin. Error bars represent 95% confidence intervals. **d.** Chemical structures of alpha-cypermethrin, a type II pyrethroid insecticide and the commonly used cytochrome P450 inhibitors, piperonyl butoxide (PBO) and clotrimazole (CTZ), shown together with the esterase inhibitor, S,S,S-tributyl phosphorotrithioate (DEF). **e.** Topology of the voltage-gated sodium channel α subunit. The four homologous repeats in the α subunit are domain-colored (from DI to DIV), each having six transmembrane segments, separated by intracellular loops (ICL); ICL1 and ICL2 and DIII-DIV linker containing MFM motif for inactivation. The positively charged S4 segments represent the voltage sensor. The pore of the channel is formed by the segments S5 and S6 and the P loops. **f.** Light photomicrograph of DUM neuron cell body isolated from the central nervous system of the cockroach *Periplaneta americana* (right) and internally perfused during whole-cell recordings of the patch-clamp technique. **g.** Representative sodium current traces, recorded according to the protocol indicated, in control and in the presence of 10 μM PBO. **h.** Comparative histogram of the sodium current amplitudes measured in control (n=27) and in the presence of 10 µM PBO (n=18), 5 µM CTZ (n=3) and 1 µM DEF (n=5), *P* values are calculated by the Mann–Whitney U test, ns, non-significant (p>0.05). Data are means ± S.E.M.; Scale bars: 40 µm and 1cm.

Inspired by these findings, we sought to determine if the limited potency of PBO-synergized pyrethroid may involve an unexpected effect of PBO on (voltage-gated sodium channels) Nav (Fig. 1e) targeted by pyrethroids^11^. We performed electrophysiological experiments (Fig. 1f) using a suitable neuronal model, the cockroach Dorsal Unpaired Median (DUM) neurons^44,45^. Intracellular application of PBO (10 µM) decreased the spontaneous sodium-dependent action potential amplitudes (Supplementary Fig. 1a-g). The effect of PBO on Nav was then confirmed in the voltage-clamp mode. PBO significantly reduced the peak current amplitudes (Fig. 1g,h). We next sought to determine whether cytochrome P450, targeted by PBO, could be specifically involved in the PBO-induced decrease of the current amplitude. We compared the current amplitudes recorded in the presence of clotrimazole (Fig. 1d), a known cytochrome P450 blocker^35,36^ and DEF (S.S.S-tributylphosphorotrithioate, Fig. 1d), an inhibitor of pyrethroid-degrading esterases^37,38^. Clotrimazole produced similar effects to those of PBO on the current amplitude, whereas DEF did not induce any effect (Fig. 1h).

To further investigate PBO-Nav interactions, we created a homology model of the cockroach PaNav1 sodium channel based on a template of the open-state structure of the rNav1.5 channel (PDB code: 7FBS) (Extended Data Fig. 1a-c). The α-subunit of the channel comprises four homologous domains, each composed of six transmembrane helices (Fig. 1e). Helices 1-4 build the voltage-sensor domains, while helices 5 and 6 of each domain contribute to the ion-conducting pore domain (the pore profile is shown in Extended Data Fig. 1b,c). A key access pathway for drugs to the center of the ion-conducting pore domain (central cavity, CC) is *via* fenestrations^39^. These four lateral tunnels connect the CC with the hydrophobic lipidic portion of the membrane (Extended Data Fig. 1d). Fenestrations are not identical (Extended Data Fig. 1e) and their diameters fluctuate during channel gating. Interestingly, we did not obtain consistent results in docking PBO to the channel. The lowest energy positions vary significantly between docking repetitions with the smina scoring function (SSF) between −4 and −6 kcal/mol (Supplementary Fig. 1h), which suggests an indirect mechanism of PBO action on Nav.

Therefore, we studied how PBO can affect the biophysical properties of Nav (Extended Data Fig. 1f-m). The voltage-dependence of the sodium conductance was shifted to more negative potentials by about 13 mV (half-activation potential) in the presence of PBO, compared to that of the control (Extended Data Fig. 1f; Extended Data Table 1; Supplementary Fig. 5a). This effect was not accompanied by a significant change in the steepness of the activation curve, parameter k (Extended Data Table 1). Neither significant differences in the voltage-dependence of inactivation (Extended Data Fig. 1g; Supplementary Fig. 5b; Extended Data Table 1) nor the use-dependent decrease in the sodium peak current amplitude (Extended Data Fig. 1h) were noted with PBO. No effect was also observed on the gaiting currents representing the outward movement of the positively charged S4 segments that initiate the opening of Nav, as shown on DUM neurons pretreated with tetrodotoxin (TTX, 50 nM) (Extended Data Fig. 1 i-m).

These data indicate that a shift to more negative potentials of activation results in an “earlier” opening of the sodium channels. The hyperpolarizing shift of activation increases the overlap of activation and inactivation (Supplementary Fig. 2a-c). This defines a range of potentials (i.e., window) where sodium channels open. PBO produced a ∼1.5-fold increase in the peak window current of the control (Supplementary Fig. 2c). At about −35 mV, the large peak-voltage of the window current predicts that sodium channels are partially inactivated, resulting in an inward current at resting membrane potential, which could produce depolarization leading to the observed decrease in action potential amplitude. Furthermore, the hyperpolarizing prepulse facilitated the recovery of only 48% of the full sodium current in the presence of PBO (Supplementary Fig. 2m).

### Deltamethrin, acting from fenestration, modifies Nav gating

To explain how PBO can impedes the effect of pyrethroids, we first reexamine the mechanisms of action of the gold-standard type II pyrethroid insecticide, deltamethrin. Figure 2a shows sodium current traces recorded in control (left) and with 1 µM deltamethrin (right), corresponding to the previously established IC ^40^. Deltamethrin significantly reduced the peak current amplitude (Fig. 2a,b). We also assessed the voltage-dependence of the conductance and the steady-state inactivation. We observed a hyperpolarized shift of the mean half-activation potential by 13.5 mV with deltamethrin (Extended Data Fig. 2a) compared to the control (Extended Data Table 1; Supplementary Fig. 5c). Unexpectedly, the voltage dependence of steady-state inactivation was strongly shifted by 21 mV toward more negative potential with deltamethrin (Extended Data Fig. 2b), compared to control (Extended Data Table 1; Supplementary Fig. 5d). In both cases, the difference in the mid-point activation and inactivation potentials occurred without any significant changes in the slope of the activation and inactivation curve parameters k (Extended Data Table 1). This indicates that the rate of inactivation from closed states was the same for Nav in control and with deltamethrin. Furthermore, the alteration in the voltage-dependence of activation and inactivation observed with deltamethrin ultimately translates into a decrease of the window current with the peak-voltage of the window shifted to the hyperpolarized potential (Supplementary Fig. 2d-f). This indicates that at voltages within this overlap region, the peak window current predicts that a smaller percentage of sodium channels will be persistently activated, compared to control and with PBO (Supplementary Fig. 2b,d,e). Moreover, the hyperpolarizing prepulses facilitated the recovery of only 43% of the full sodium current, suggesting that recovery from fast inactivation was also affected by deltamethrin (Supplementary Fig. 2n). We also examined the use-dependent action of 1 µM deltamethrin (Extended Data Fig. 2c). Upon multiple depolarizations, the control peak sodium current amplitude, recorded within the pulse, decreased with the number of pulses to reach a steady-state level after 10 pulses. The use-dependent decrease in the sodium peak current amplitude was stronger in the presence of deltamethrin, with significant decreases in peak current after 10 pulses. Collectively, these results indicate that deltamethrin-induced changes in the voltage dependence of both activation and inactivation towards hyperpolarized potentials may lead to neuronal hyperexcitability, allowing the deltamethrin use-dependent effect. These hyperpolarized shifts tend to reduce the window current and thus, the fraction of permanently activated Nav channels (i.e., decrease in current amplitude). Therefore, these results show that deltamethrin interacts with the open state but also stabilizes the fast inactivated state of Nav.

**Fig. 2.**
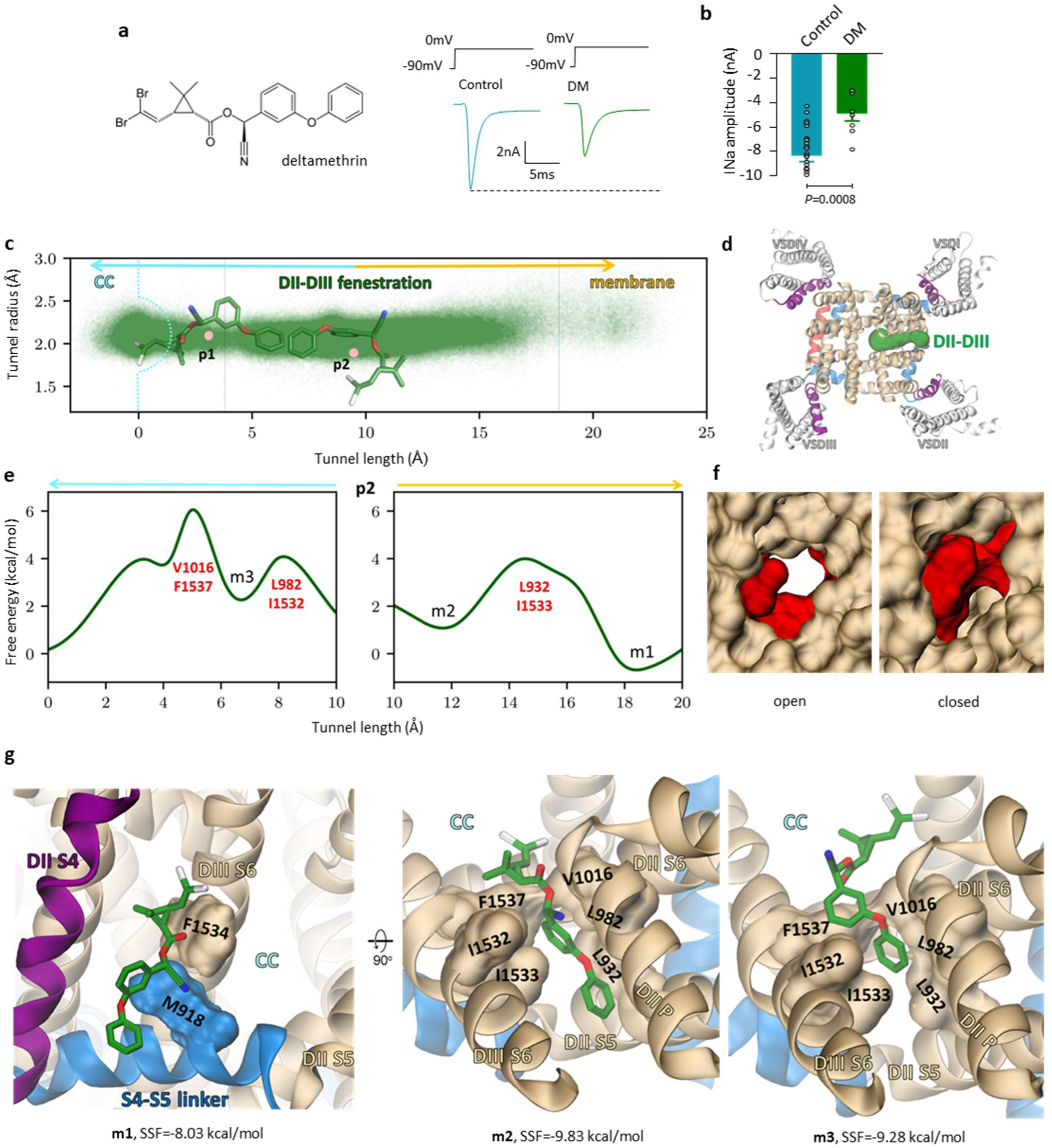
Deltamethrin acts through the DII-DIII fenestration. **a.** Representative sodium current in control and in the presence of 1 μM deltamethrin (DM, chemical structure in *inset*). **b.** Histogram summarizing the effect of DM (1µM, n=8) on sodium current amplitude (control, n=27). Data are means ± S.E.M.; *P* values are calculated by the Mann–Whitney test. **c.** Deltamethrin positions in the DII-DIII fenestration, shown as green dots representing the geometric center (GC) of the ligand, sampled during 30 MD trajectories along the tunnel. The GCs of deep (p1) and shallow (p2) docking poses of the ligand are represented by pink dots. The tunnel begins in the central cavity (CC) of the channel and ends at the plasma membrane interface. Thin vertical lines show bottlenecks caused by two free energy barriers. **d.** The DII-DIII fenestration tunnel is shown schematically in the top view of the PaNav1 model. **e.** Free energy profiles calculated by well-tempered metadynamics initiated from the shallow pose (p2) toward the CC (left) and the fenestration entry (right). Bottleneck residues in the tunnel between energy minima (m1-m3) are marked in red. They are also shown in the surface representation of the DII-DIII fenestration of both open and closed channel conformations in (**f**). Ligand poses corresponding to the energy minima m1, m2, and m3 are shown in (**g**). For the m2 and m3 minima, bottleneck residues are shown in the surface representation. Binding energies are calculated using smina scoring function (SSF). Residues are renumbered based on the house fly sodium channel (*Musca domestica*, GenBank accession number: X96668).

We then proceed to find deltamethrin binding sites on the Nav channel. As pyrethroids are known to bind to the open-state channels, we began by docking to the PaNav1 model in the open state. We obtained two clusters of poses in the DII-DIII fenestration: (i) deeply bound in the tunnel, with the halogen part reaching the CC of a channel (p1, Fig. 2c), and (ii) shallow binding at the membrane-fenestration interface (p2, Fig. 2c). This observation led us to question whether fluctuations of the fenestration diameter (Fig. 2d) enable deltamethrin to bind deeply in the tunnel, where it can reach the residues frequently found mutated in pyrethroid-resistant insects by a mechanism called knockdown resistance (kdr). Such interaction could not be explained by any docking studies presented before. We thus used well-tempered metadynamics to investigate the deltamethrin binding pathways along the fenestration. The reconstructed free-energy profiles revealed three distinct free-energy minima (m1-m3, Fig. 2e) occupied by the ligand. The m1 minimum is located at the channel-membrane interface where deltamethrin interacts with the M918 and F1534, frequently found mutated in the resistant insects. In the m2 and m3 minima, closely related to p2 and p1 identified by docking (Fig. 2c), deltamethrin penetrates the fenestration and interacts with multiple residues from DII and DIII pore domain helices (Extended Data Fig. 3a-d). Mutations in those positions affect pyrethroid action on Nav (for the list of kdr residues interacting with deltamethrin, see Extended Data Table 2). Moreover, ligand bound in the m3 minimum interacts with V410 and N1013, residues contributing to the second pyrethroid receptor site (PyR) proposed before (Extended Data Fig. 3c,d), thus linking two PyRs into one. Pairs of fenestration bottlenecking residues (marked red in Fig. 2e,f) create free-energy barriers limiting the mobility of the ligand. While widespread substitutions of lysines to the bulkier aromatic residues (L932F, L982W) reduce pyrethroids efficacy on Nav^15,41–44^ and tunnel diameter, the I1533A substitution enhances channel sensitivity to deltamethrin^45^, suggesting a stronger effect of pyrethroids when deeply bound in the fenestration. Finally, the residues creating the third free-energy barrier and the narrowest point in the fenestration, V1016 and F1537, contribute most to the deltamethrin binding. They are crucial in ligand interaction with the channel in both the m2 and m3 minima and block full entrance to the CC. V1016I/G is one of the most frequently reported kdr mutations worldwide, while the F1537A substitution (F1518A in BgNav) was found to diminish channel sensitivity to deltamethrin and other pyrethroids examined^45^. The deltamethrin-channel affinity is stable when the ligand crosses the fenestration (Extended Data Fig. 3e).

The comparison of the DII-DIII tunnel profiles between open, inactivated, and closed static models (Extended Data Fig. 3f) indicates that conformational changes of the ion-conducting pore are coupled with changes in the dimensions of fenestrations. While the DII-DIII profile is similar in the open and inactivated states, the tunnel of the closed model is much narrower. Moreover, the fluctuations of the bottleneck radius (BR) observed in classical molecular dynamics (MD) simulations are prominent in the inactivated state of Nav where a bimodal distribution of BR was found. In contrast, the average value of BR equals 1.3 Å in the Nav open state (Extended Data Fig. 3g). This made us wonder whether binding to the inactivated state Nav could also be possible.

Thus, we docked deltamethrin to the PaNav1 models representing inactivated and closed states to investigate the state dependence of binding. The lowest energy pose in the inactivated state was found in the same region as the deeply bound pose in the open model with the backbone root-mean-square deviation (RMSD) equal to 3.38 Å (Extended Data Fig. 4a). In both models, deltamethrin forms π-stacking interactions with F1537 and the α-cyano groups of the ligand overlap. No binding poses were found in the fenestration of the closed model. The alignment of these three models shows that there is a shift of the fenestration-bottlenecking F1537 (RMSD=0.21 Å between the open- and inactivated-state and RMSD=1.42 Å between the open- and closed-state models), which precludes deltamethrin binding to the closed state Nav due to a steric clash (Extended Data Fig. 4b). Interestingly, in one of our 500 ns-long MD trajectories starting from the open state Nav, we found that the channel spontaneously inactivates (the MFM motif of the inactivation particle moves to its binding pocket). We excluded this trajectory from fenestrations analyses as an outlier but we took advantage of the opportunity to investigate the possibility of deltamethrin binding when the channel undergoes a conformational transition. We docked deltamethrin to frames extracted from this simulation, having the binding site defined based on previously found “deep” position in DII-DIII fenestration. The smina scoring function (SSF) fluctuated around −7.85 kcal/mol with the SD=0.58 kcal/mol (Extended Data Fig. 4c). These results indicate that deltamethrin can enter the DII-DIII fenestration when the channel is in the open state and that the interaction can last when the channel undergoes a conformational transition to the inactivated state.

### PBO impedes deltamethrin effect *via* the cytochrome P450

We then focused on how PBO could alter deltamethrin efficacy. After intracellular dialysis with 10 µM PBO, the fraction of the remaining residual sodium current was not further reduced when 1 µM deltamethrin was applied (Fig. 3a). The voltage-dependence of steady-state inactivation was shifted by about 10 mV toward more positive potentials compared to DUM neurons treated with deltamethrin alone. By contrast, the voltage-dependence of the conductance was not statistically changed (Extended Data Fig. 2d,e; Extended Data Table 1). The superimposed activation and steady-state inactivation curves revealed a ∼3-fold increase in the hyperpolarized shifted peak-voltage window current compared to that of with deltamethrin alone (Supplementary Fig. 2h,i). In this case, PBO impedes the effect of deltamethrin, as described above, by shifting the inactivation to more positive potentials, thereby expanding the window current.

**Fig. 3.**
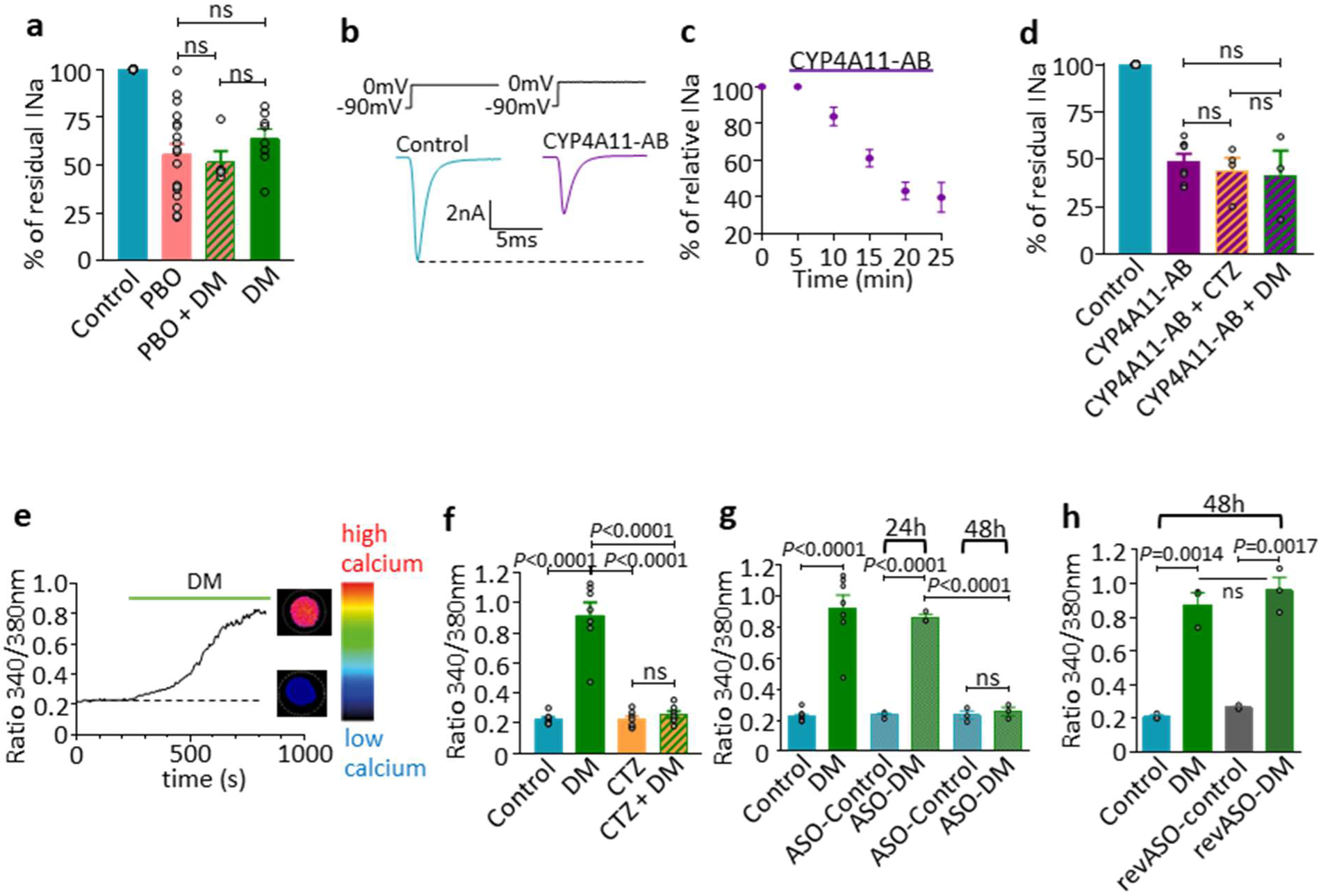
Inhibition of CYP4 hampers the action of deltamethrin on voltage-gated sodium channels. **a.** Histogram illustrating the residual sodium current amplitude (%) recorded in control (n=27), with 1 µM DM alone (n=8), in the presence of 10 µM PBO, applied alone (n=18) and with 1µM DM (n=3). **b.** Sodium currents recorded just after forming the whole-cell configuration (control) and 25 minutes following the dialysis with 5 µg/mL CYP411 antibody (CYP411-AB). **c.** Relative peak sodium current (%) plotted *versus* different times, following the dialysis with 5 µg/mL CYP411-AB. Time zero was marked at the time of rupture of the membrane patch for whole-cell recordings (n=4-10). **d.** Residual sodium current amplitude (%) recorded under intracellular application of 5 µg/mL CYP411-AB alone (n=8), with 5 µM clotrimazole (CTZ, n=4) and 1 µM DM (n=3); ns, non-significant (p≥0.05). **e.** Progressive [Ca^2+^]_i_ rise in Fura-2-loaded DUM neuron (*Inset*) plotted *versus* time of application of 1 µM DM. **f.** Histogram showing the [Ca^2+^]_i_ rise recorded in control (n=7), with DM (1 µM) alone (n=7), with CTZ (clotrimazole, the inhibitor of cytochrome P450 enzyme) alone (5 µM, n=7) and with DM (1 µM) aftter pretreatment with CTZ (5 µM, n=7). **g.** Knockdown of CYP4 by antisense oligonucleotides (ASO) inhibited the DM-induced elevation of [Ca^2+^]_i_ (n=7). DUM neurons were incubated with 10 µM ASO for 24 h and 48 h alone (n = 3) and after application of 1 µM DM (n=3), ns, non significant (p≥0.05). **h.** Histogram illustrating the lack of effect of reverse antisense oligonucleotides (revASO, 10 µM) after 48 h, applied alone (n=3) and following application of 1 µM DM (n=3). **a,c,d,f,g,h,** Data are means ± S.E.M.. **f,g,h** *P* values are calculated by the Student unpaired t-test **a,d,** *P* values are calculated by the Mann–Whitney U test, ns, non-significant (p≥0.05).

Next, we studied the involvement of cytochrome P450 enzymes in the modulation of the effect of deltamethrin on Nav. In the American cockroach *Periplaneta americana*, genomics analyses reveal expansion in the gene families encoding for detoxification enzymes. Cytochrome P450 clustered particularly with the lineages of CYP4 genes^46,47^. We internally dialyzed DUM neurons with cytochrome P450 polyclonal antibody CYP411, which binds to insect CYP4. The sodium current amplitude decreased within 5 to 25 min after intracellular dialysis with the antibody (5 µg/mL) (Fig. 3b-d). The fraction of the residual sodium current remaining after the application of CYP411 was not further reduced by neither 5 µM clotrimazole (a cytochrome P450 inhibitor) nor 1 µM deltamethrin (Fig. 3d). These findings indicate that cytochrome P450 inhibition contributes to modulating the efficacy of deltamethrin. Deltamethrin also increases the intracellular calcium concentration [Ca^2+^]_i_ in insect neurons through Nav and the reverse Na/Ca exchanger^48^. We then examined whether the deltamethrin-induced elevation of [Ca^2+^]_i_ (Fig. 3e) was affected by clotrimazole. Pretreatment of DUM neurons with clotrimazole (5 µM) completely abolished the effect of deltamethrin (Fig. 3f). We also examined the effects of antisense oligonucleotides (ASO) directed against CYP4. Based on data obtained with the CYP4A11 antibody, we designed ASO (Supplementary Table 1) to decrease mRNA expression of CYP4 family of cytochrome P450. DUM neurons were treated for 24h or 48h with ASO (Fig. 3g). As shown, CYP4 ASO completely abolished [Ca^2+^]_i_ rise induced by deltamethrin after 48h pretreatment. By contrast, deltamethrin still increased [Ca^2+^]_i_ in DUM neurons treated for 48h with reverse antisense oligonucleotides (Fig. 3h; Supplementary Table 1), which excludes unspecific effects. These results demonstrate that mRNA expression of the CYP4 family of cytochrome P450 was suppressed in DUM neurons pretreated by ASO and confirm that cytochrome P450 plays substantial roles in the reduction of deltamethrin effects.

Next, we used immunocytochemistry to determine whether Nav and cytochrome P450 CYP4 were localized in the same cellular region to envisage potential interactions of these two proteins. Images captured with confocal laser scanning microscopy show positions of each single label, revealed using their specific excitation spectra (Extended Data Fig. 5b,c and e,f). Anti-SP19 antibodies are directed against the domain III-IV linker containing the MFM motif of the inactivation particle of Nav-α subunit. These antibodies are commonly used to reveal the expression of Nav in insects and specifically in DUM neurons^49,50^. SP19-immunoreactivity was most intense close to the initial segment, which is in agreement with previous SP19 immunostaining^49^. Regarding the CYP4A11 immunoreactivity, the intensity of the staining represented by small points was also preferentially localized at the basal pole close to the initial segment showing a clear overlap with the distribution of SP19 immunoreactivity (Extended Data Fig. 5a-c; d-f). These results indicate a spatial overlap close to the initial segment between two different fluorescent labels, FITC and Alexa-Fluor 633, with specific primary antibodies directed against Nav and the cytochrome P450 CYP4 (Supplementary Fig. 3a-c; d-f).

### Octopamine regulates the cytochrome P450-induced modulation of Nav

In insects, the stress reaction is related to the release of the biogenic amine octopamine, involved in the regulation of diverse physiological mechanisms ^32,51^. Hence, we investigated whether octopamine regulates cytochrome P450 and thus deltamethrin. Using immunocytochemistry and calcium imaging, we showed that octopamine (1 µM) increased intracellular levels of the second messenger cAMP^52^ and [Ca^2+^]_i_ (Supplementary Fig. 4a-c). These effects were completely blocked by the standard octopamine receptor antagonist, phentolamine, and a selective α-adrenergic-like octopamine receptors antagonist, chlorpromazine^53^ (Supplementary Fig. 4b,c). Octopamine receptors expressed by DUM neurons might thus be more closely related to α-adrenergic-like octopamine receptor, confirming previous cloning data obtained in *Periplaneta americana*^54^.

In addition, we previously documented the involvement of DARPP-32 (dopamine- and cAMP-regulated neuronal phosphoprotein-32)^55^ in both calcium and cAMP signalings following activation of DUM neuron octopamine receptors^52^ through G proteins, involving Gαq and Gαs subunits, respectively (Fig. 4a). To assess the role of DARPP-32 in regulating deltamethrin efficacy, we specifically inhibited/activated the different intracellular cascades involved in the DARPP-32 phosphorylation (*via* protein kinase A, PKA) or dephosphorylation (*via* protein phosphatase PP2B)^52^. To mimic the effect of octopamine (1 µM) *via* Gαs protein activation, we used 8-Br-cAMP. We observed a decrease of the [Ca^2+^]_i_ rise produced by deltamethrin. By contrast, this effect was not observed in the presence of the PKA inhibitor, H89 (Fig. 4a,b, blue pathway). Intracellular application of the PP2B inhibitor, cyclosporine A, completely blocked the deltamethrin-induced elevation of [Ca^2+^]_i_, when PKA was inhibited by H89 (Fig. 4a,c; green pathway). Moreover, we previously showed that the phosphorylated DARPP-32 is a potent inhibitor of the protein phosphatase PP1/2A^52^. Okadaic acid (OkAC), an inhibitor of PP1/2A, tested in the presence of H89, fully inhibited the deltamethrin-induced elevation in [Ca^2+^]_i_ (Fig. 4a,d). These results indicate that DARPP-32 phosphorylated by PKA inhibits the phosphatase PP1/2A. The resulting inhibition of the phosphatase mimics the effect of cytochrome P450 inhibition (Fig. 4a), which strongly reduces deltamethrin efficacy on Nav. DARPP-32 phosphorylation is opposed by a dephosphorylation process that involves the calcium/calmodulin sensitive phosphatase PP2B, following activation of Gαq protein and the consecutive [Ca^2+^]_i_ rise^52^ (Fig. 4a). In this case, the inhibition of phosphatase PP1/2A was completely reversed (Fig. 4d), restoring the full effect of deltamethrin.

**Fig. 4.**
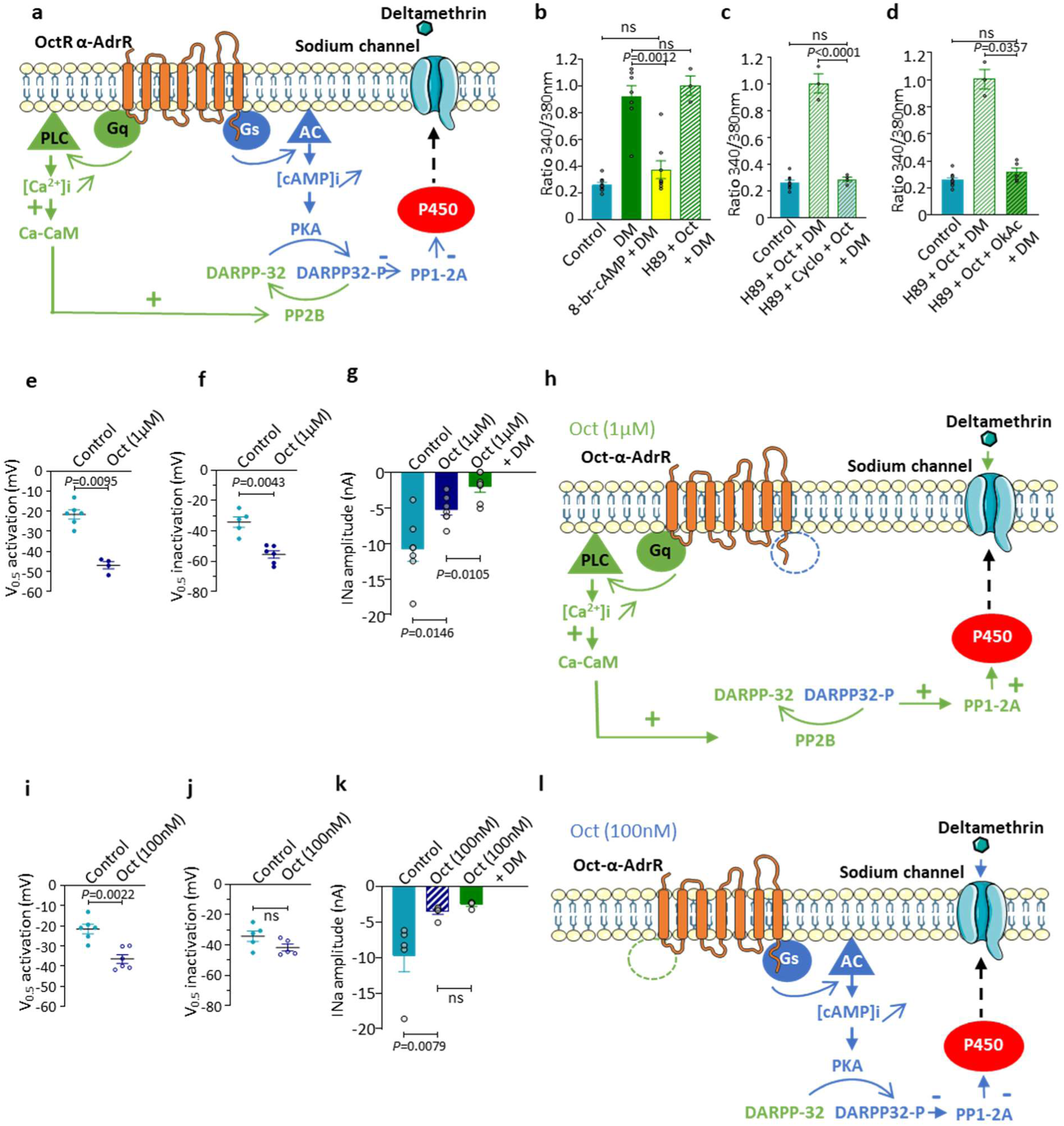
Inhibition of CYP4 hampering deltamethrin effect requires the α-adrenergic-like octopamine receptor signal pathway in DUM neurons. **a.** Summarizing molecular model of mechanisms of DUM neuron-expressed α-like adrenergic receptor (OctR α-AdrR) involved in the CYP4 blockade and the inhibition of deltamethrin action on sodium channel through DARPP-32 (Dopamine- and cAMP-Regulated Phosphoprotein, Mr=32 KDa) and DARPP-32-phosphorylation process. **b-d.** Comparative histograms illustrating the Fura-2 fluorescence ratio (340/380) of [Ca^2+^]_i_ recorded under different experimental conditions indicated below each bar. Control (n=9), DM, (1 µM deltamethrin, n=7); 8-Br-cAMP (1 µM) +DM (1 µM), (n=8); H89 (PKA inhibitor), (1 µM) + Oct (octopamine) (1 µM) + DM (1 µM), (n=3); H89 (1 µM) + Cyclo (cyclosporine A, PP2B inhibitor) (100 nM) + Oct (1 µM) + DM (1 µM) (n=4); H89 (1 µM) + Oct (1 µM) + OkAc (okadaic acid, PP1-2A inhibitor) (1 µM) + DM (1 µM) (n=5). Data are means ± S.E.M.. **b-d,** *P* values are calculated by the Mann–Whitney U test. ns, non-significant (p≥0.05). **e.f.** Scatter-plots illustrating the values of mid-point (V_0.5_) of activation (**e**) and inactivation (**f**) for sodium current in control (n=6 and n=5, respectively) and in the presence of higher concentration of octopamine (Oct, 1 µM) (n=4 and n=6, respectively). **g.** Comparative histogram illustrating the sodium current amplitude, in control (n=7), with Oct (1 µM, n=7) and with Oct (1 µM) mixed with DM (1 µM, n=7). Data are means ± S.E.M.. *P* values are calculated by the Mann–Whitney U test. **h.** Hypothetic model illustrating the specific implication of Gαq-coupled Oct-α-AdrR leading to the dephosphorylation of DARPP-32-P to DARPP-32 through calcium/calmodulin-dependent protein phosphatase 2B (PP2B). This results in the activation of cytochrome P450. **i,j.** Scatter-plots showing the values of half-maximal voltage-dependence (V_0.5_) of activation (**i**) and inactivation (**j**) for sodium current in control (n=6 and n=5, respectively) and in the presence of low concentration of octopamine (Oct, 100 nM) (n=6 and n=5, respectively). **k.** Comparative histogram representing the sodium current amplitude recorded at 0 mV from a holding potential of −90 mV in control (n=5), with a low concentration of Oct (100 nM, n=5) and with Oct (100 nM) mixed with 1 µM DM (n=4). Data are means ± S.E.M.. *P* values are calculated by the Mann–Whitney U test. ns, non-significant (p≥0.05). **l.** Hypothetic pattern summarizing the essential intracellular molecular events involved in the inhibition of cytochrome P450 through Gαs-coupled receptors (Oct-α-AdrR), cAMP/PKA cascade and phosphorylated DARPP-32, activated by lower concentration of octopamine.

Given these findings, we tested whether octopamine, used at different concentrations, could specifically modulate the effect of deltamethrin through activation of octopamine receptors coupled to Gαs and Gαq subunits mediating elevation of [cAMP]_i_ and [Ca^2+^]_i_ respectively. Octopamine at 100 nM and 1 µM reduced sodium current amplitudes (Fig. 4g,k), as it previously documented elsewhere^34^. Octopamine (1 µM) produced a hyperpolarized shift of both voltage-dependent conductance and steady-state inactivation (Fig. 4e,f; Extended DataTable 1), whereas octopamine at 100 nM only shifted the voltage-dependence of the conductance toward more negative potentials (Fig. 4i,j; Extended Data Table 1). These concentration-dependent effects of octopamine were statistically significant (Supplementary Fig. 6g,h). The most interesting result was that at 1 µM octopamine, the sodium current amplitude was further reduced in the presence of deltamethrin (Fig. 4g), as it was observed when DARPP-32 was dephosphorylated (Fig. 4h, green pathway). The superimposed conductance and steady-state inactivation curves revealed a pronounced hyperpolarized shift of the peak-voltage window current (Supplementary Fig. 6a-c and d-f). These results confirm the hypothesis that the effect of deltamethrin on Nav is favored when the inactivated state is stabilized. They also suggest that octopamine (1 µM) preferentially activates signaling cascade through Gαq coupling rather than Gαs (Fig. 4h).

We performed additional experiments with 100 nM octopamine, which corresponds closely to the level of octopamine detected under stressful conditions^56^. We previously showed that at 100 nM, octopamine only shifted the voltage-dependence of the conductance toward more negative potentials (Fig. 4i,j; Extended Data Table 1). But in contrast to octopamine used at 1 µM, non-significant additional effect of deltamethrin on the sodium current amplitude was observed (Fig. 4k). Based on the previous findings (Fig. 4a-d), this lack of effect of deltamethrin seems to be mediated *via* Gαs-coupled receptors and cAMP/PKA cascade (Fig. 4k,i). Furthermore, these results also reinforce the hypothesis that the effect of deltamethrin on Nav is favored when its inactivated state is stabilized. This was confirmed by experiments indicating that, in the presence of 100 nM octopamine, deltamethrin fully inhibited the sodium current amplitude when a depolarizing pulse was evoked from a holding potential of −50 mV compared to −90 mV (Supplementary Fig. 6i,j).

### DARPP-32 modulates deltamethrin efficacy on Nav

Although high concentration of octopamine (i.e., 1 µM) is not so physiological^56^, the opposite effect reported above allow us to use this alternative approach to further elucidate the regulatory mechanisms involved in the modulation of the effect of deltamethrin *via* cytochrome P450. We then tested the effect of octopamine (1 µM) on the sodium current amplitude while maintaining the cytochrome P450 enzymes inhibited by PBO. According to this protocol (Extended Data Fig. 6a1), octopamine at 1 µM reduced the current amplitude in DUM neuron dialyzed with PBO (10 µM). More interestingly, even in the presence of PBO, deltamethrin (1 µM) reduced the current amplitude under octopamine (1 µM) treatment (Extended Data Fig. 6b,1c). Knowing that PP2B-induced dephosphorylation of DARPP-32, which thereby reverses the inhibition of PP1/2A and cytochrome P450, we then tested okadaic acid, previously used to inhibit PP1/2A. We internally dialyzed DUM neurons with PBO (10 µM) and okadaic acid (1 µM) (Extended Data Fig. 6a2). After 20 minutes, treatments with octopamine (1 µM) alone and with octopamine mixed with deltamethrin (1 µM) were studied. In this case, deltamethrin did not produce any effect on sodium current amplitude (Extended Data Fig. 6b2,c). In addition, the hyperpolarized shift of both voltage-dependence of conductance and steady-state inactivation was observed with PBO/octopamine/deltamethrin when compared to PBO/deltamethrin (Extended Data Fig. 6d). This was completely reversed in the presence of okadaic acid (Extended Data Fig. 6e,f; Extended Data Table 1). These results underscore an unexpected mechanism by which PP1/2A regulated by DARPP-32/DARPP-32-P was involved in the modulation of cytochrome P450 function leading to decrease or increase deltamethrin efficacy on Nav, respectively. Finally, the significant effect of deltamethrin on Nav under octopamine (1 µM) observed in the presence of PBO, suggests that activation of PP1/2A might change the conformation-activity relationships of cytochrome P450. This could modify the shape of the malleable and flexible active-site occupancy of enzymes^57^, which alters its stereoselectivity for PBO. The resulting decrease of PBO affinity could render heme cofactor available, which is suspected to be involved in the indirect interaction with Nav and the effect of deltamethrin.

### Intracellular hemin alters Nav and increases deltamethrin efficacy

Heme is a cofactor essential for cytochrome P450 enzymes. Recent data indicate that heme and its oxidized form, hemin, play important roles as voltage-gated ion channel modulators^58^. We thus tested the effects of hemin on the sodium current amplitude and the gating properties of Nav (Extended Data Fig. 7a). Intracellular application of hemin (100 nM) significantly reduced the peak sodium current amplitudes (Extended Data Fig. 7b). In addition, the sodium current amplitude decreased over periods of 3 to 10 min after intracellular dialysis of hemin before reaching a steady-state level (Extended Data Fig. 7c). Moreover, we observed a hyperpolarized shift of the mean half-activation potential by about 13 mV with hemin compared to control (Extended Data Table 1; Extended Data Fig. 7d; Supplementary Fig. 5e). The voltage-dependence of steady-state inactivation was also shifted by about 15 mV toward more negative potential with hemin, compared to control (Extended Data Table 1; Extended Data Fig. 7e; Supplementary Fig. 5f). This suggests that hemin could also interact with the inactivated state of Nav. Furthermore, the alteration of the voltage-dependence of activation and inactivation observed with hemin translates into a decrease of the window current with the peak-voltage of the window shifted to the hyperpolarized potential (Supplementary Fig. 2j-l). This indicates that at voltages within this overlap region, a smaller percentage of sodium channels will be persistently activated, as compared to the control. Very similar results were obtained with deltamethrin (Supplementary Fig. 2d-f). Furthermore, the hyperpolarizing prepulses facilitated recovery of only 32% of the full sodium current, suggesting that recovery from fast inactivation was also affected by hemin (Supplementary Fig. 2o). To test for the specificity of the hemin effect, we measured the impact of its component, Fe^2+^, using FeSO_4_. The sodium current amplitude was unaffected after the application of FeSO_4_ (1 µM) (Extended Data Fig. 7f).

We then tested whether hemin modified the effect of deltamethrin. As expected, the fraction of the residual sodium current remaining after application of hemin was further reduced by deltamethrin (1 µM) (Extended Data Fig. 7g). Finally, we also examined the effect of intracellularly applied deferoxamine (DFO), the iron chelator, known to inhibit heme/hemin effects^59,60^ (Extended Data Fig. 7h). Similar to PBO, the fraction of the remaining residual sodium current observed after the application of DFO (10 µM) was not further reduced when 1 µM deltamethrin was co-applied with DFO (Extended Data Fig. 7i). Furthermore, deltamethrin did not decrease the sodium current amplitude when tested with octopamine at 1 µM and DFO. This suggests that DFO, a potent iron chelator, could form a stable complex with iron of cytochrome P450 that affects the deltamethrin effect (Extended Data Fig. 7j).

These results suggest that heme iron centers could represent an important element for regulating Nav function and deltamethrin efficacy. It is known that heme iron provides positively charged residues. The modification of the shape of the cytochrome P450 enzyme active site, suggested above, can introduce new repulsive forces between cytochrome P450 and relevant segments of Nav, modulating the channel function. We thus performed molecular modeling analysis to investigate the influence of the electrostatic surface potential of the intracellular side of sodium channel PaNav1 depending on the main conformationally distinct states. The position of the DIII-DIV linker differs greatly between the models representing open, inactivated, and closed states (Fig. 5a). The RMSD of the inactivation particle motif MFM is 13 Å for the open, and 18.9 Å for the closed model when compared to the inactivated state. In the closed model, the DIII-DIV linker is shifted towards the channel axis while being deeper in the cytosol. The electrostatic potential surfaces show that the positive charges are exposed to the cytosol in the inactivated (Fig. 5b) and open states, while in the closed model, five lysines face the transmembrane part of the channel (Fig. 5c).

**Fig. 5.**
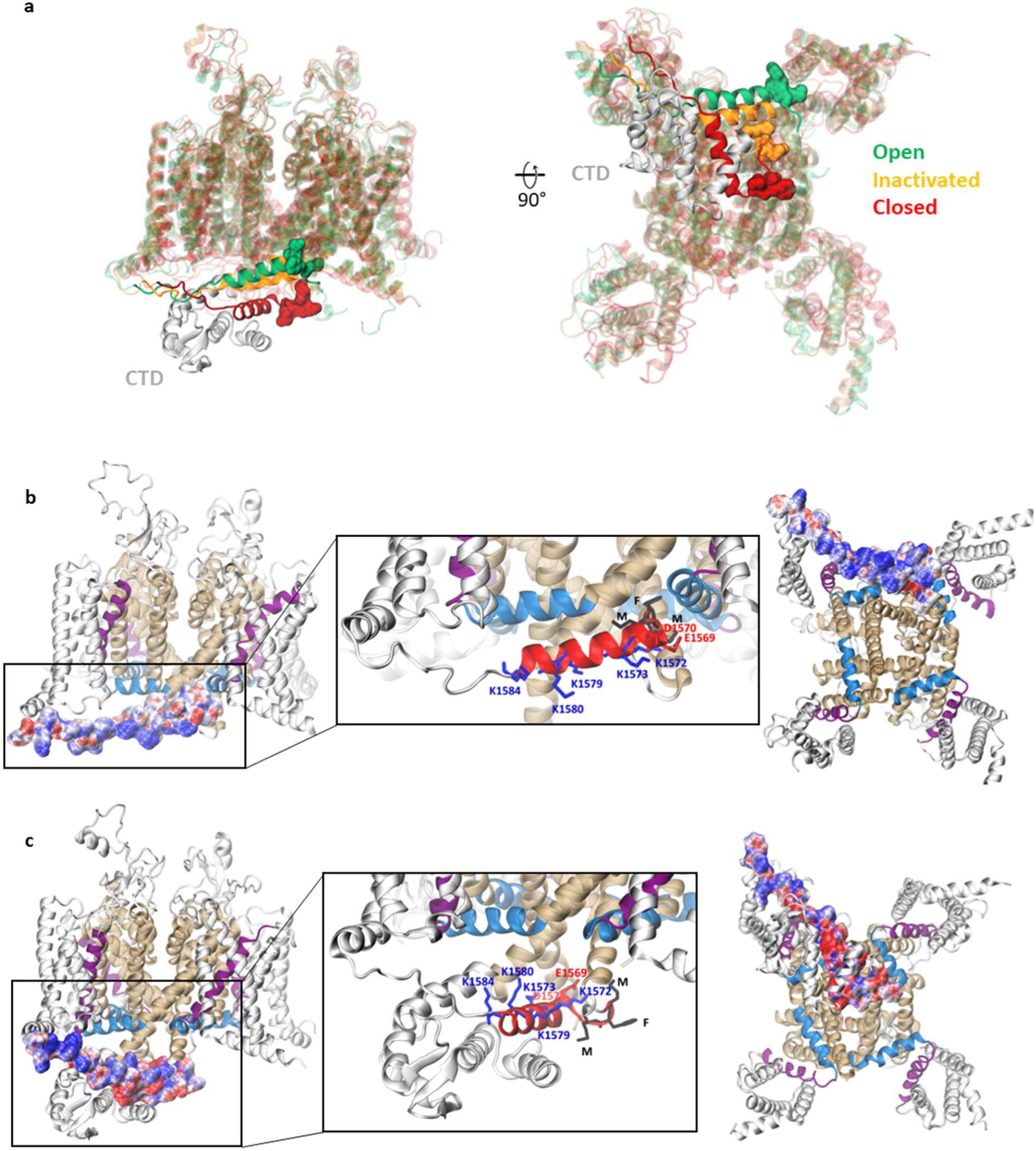
Overview of the 3D structure of the sodium channel PaNav1 with relevant segments, residues, and electrostatic surface potential. **a.** Alignment of PaNav1 models in the open (green), inactivated (orange), and closed (red) states in the side (left panel) and bottom (right panel) view. The DIII-DIV linker is highlighted and its inactivation particle MFM motif is shown in the surface representation. The carboxy-terminal domain (CTD) is present only in the closed state model. **b,c.** The electrostatic surface potential of the DIII-DIV linker is mapped on the PaNav1 in the inactivated (**b**) and closed (**c**) states. Side views are shown on the left panels, and bottom views are shown on the right panels. In the middle panel, charged amino acid residues are shown in sticks (blue – positively charged, red – negatively charged), and the MFM motif of inactivation particle is shown in black.

### Chemical stress reduces deltamethrin efficacy

To assess the physiological relevance of our results, we performed toxicological experiments *in vivo* on the cockroach *Periplaneta americana*. We exposed cockroaches for 45 minutes to low concentration of the chemical stressor, the insecticide bendiocarb, and evaluated the octopamine level using LC-MS/MS. In control insects, the octopamine level was equal to 56.58 ± 5.05 ng/g, while after 0.1 nM bendiocarb exposure it increased to 84.72 ± 12.06 ng/g (p=0.04) (Extended Data. Fig. 8a,b). We then evaluated the effectiveness of deltamethrin in insects pre-exposed and not exposed to 0.1 nM bendiocarb. In the cumulative survival analysis, the median survival time for insects subjected to deltamethrin was 31.4 hours, while in insects pre-exposed to 0.1 nM bendiocarb, this time was extended to 34.98 hours (Extended Data Fig. 8c). A clear shift in the survival curve observed in insects pre-exposed to bendiocarb indicates a lower deltamethrin effectiveness in this group. As survival analysis required higher doses of neuroactive compound, we also evaluated the potency of deltamethrin in the insect motor ability test (time of turning back from dorsal to ventral side), where the first signs of paralysis can be observed at smaller concentrations. Control insects turned back in 1.65 ± 0.36 s, while insects treated with 1 µM deltamethrin for 1 hour had visible symptoms of paralysis and their time of turning back increased to 6.59 ± 1.85s (p=0.01). In a group of insects pre-exposed to 0.1 nM bendiocarb, the time of turning back was similar to the control level (1.61 ± 0.30s; p=0.02 in comparison to insect treated with 1 µM deltamethrin) (Extended Data Fig. 8d). Overall, these results indicate that the exposition to the chemical stressor, bendiocarb, increases the octopamine level, which further reduces deltamethrin efficacy.

## Discussion

Piperonyl butoxide is used worldwide in formulations with pyrethroids, the first line insecticides, to limit metabolic detoxification by inhibiting the cytochrome P450. However, recent studies raise questions about the unexpected weak performance of PBO in enhancing the pyrethroid efficacy without providing explanation thereof. Here, we demonstrate that the PBO-induced inhibition of cytochrome P450 impedes the effect of deltamethrin by mainly affecting its interaction with the inactivated state of Nav. Our findings provide insights into a new mechanism underscoring the cytochrome P450-induced regulation of Nav. The mechanism involves DARPP-32, following activation of α-adrenergic-like octopamine receptors, which in turn regulate cytochrome P450. These findings reveal a new adaptation mechanism developed by insects to reduce the efficacy of insecticides.

We started by refining our understanding of the deltamethrin mode of action on Nav. Deltamethrin induced changes in the voltage-dependence of both activation and inactivation towards hyperpolarized potentials. Together with the use-dependent of its action, the results indicate that deltamethrin affects both open and inactivated state of Nav. By molecular docking followed by enhanced sampling simulations, we showed how deltamethrin, acting from the DII-DIII fenestration, interacts with multiple residues scattered on Nav protein that confer resistance to pyrethroids when mutated. By analyzing the free-energy profiles of the deltamethrin pathway through the fenestration, we described the role of kdr mutations that increase or decrease the fenestration diameter, thus increasing or decreasing the probability of deltamethrin reaching its final binding site near the ion-conducting pore. The bottleneck radius of the DII-DIII fenestration is wider in the inactivated than in the open state, suggesting that deep binding is favored when the channel inactivates. Based on docking results to the static Nav models representing closed, open, and inactivated states, we suggest that our free energy minima correspond to binding to a given functional state of the channel: m1 – closed, m2 – open, and m3 – late open and inactivated. This hypothesis is strengthened by deltamethrin docking to the Nav that spontaneously inactivates, where the SSF is roughly constant. The interaction with the inactivated state, not described before, may help to understand why PBO impedes deltamethrin action on Nav.

Our findings strongly suggest an indirect mechanism of PBO action on Nav through the involvement of cytochrome P450. These enzymes are hemoproteins^20,61^ in which the heme iron is typically ferric (Fe^3+^) in the resting state but can be ferrous (Fe^2+^) *in vivo*^62^. It has been recently reported that heme and its oxidized form hemin (Fe^3+^) are non-genomic modulators of different voltage-gated ion channel functions^58,63,64^. In this context, we showed that intracellularly applied hemin affects Nav similarly to deltamethrin. The voltage-dependence shift of activation and inactivation parameters suggests that hemin can modulate the open state but its action is favored when the fast-inactivated state is more preponderant. It is thus not surprising that after pretreatment with hemin, the effect of deltamethrin was more pronounced. Such potentiation of deltamethrin action was not observed with the iron chelator, DFO^59,60^, which probably binds to heme/hemin *via* the iron moiety. Although it remained unclear how the iron-containing porphyrin could act as a gating modifier of Nav, an interesting hypothesis emerged from our molecular modeling comparison of the cockroach Nav (PaNav1), representing the three primary states: closed, activated, and inactivated. The electrostatic potential surfaces of the intracellular DIII-DIV linker, containing the inactivation particle, show that the positive charges are exposed to the cytosol in the open and the inactivated states but not in the closed state. From these results, together with electrophysiological data, it is tempting to speculate that the strength of charge–charge electrostatic repulsion between the iron-containing porphyrin and the positive charges exposed to the cytosol in the inactivated state may affect the kinetic properties of Nav by stabilizing the fast-inactivated state. Although the heme of cytochromes P450 is not directly accessible, it has previously been reported that the active sites differ considerably in flexibility and susceptibility. The overall shape of the active-site cavity containing the heme is accompanied by large changes in the size of the active site^57,61,65–67^. This gives us information on how cytochromes with different folds i) can interfere with the heme properties^68,69^ and ii) could facilitate electrostatic interactions with positively charged residues exposed to the cytosol in the inactivated state of Nav. These conformational dynamics and/or plasticity of cytochrome P450 structure might be achieved by a phosphorylation/dephosphorylation process (e.g., cAMP-dependent PKA)^70,71^. Here, we demonstrate a sophisticated regulation of cytochrome P450 by phosphorylation/dephosphorylation involving octopamine, DARPP-32 and PP1/2A (Figure 4).

We show that a high octopamine concentration (1 µM) acting through Gαq activated pathway and PP2B, produces dephosphorylation of DARPP-32. The resulting activation of PP1/2A removes cytochrome P450 inhibition potentiating the effect of deltamethrin. Octopamine (1µM), like deltamethrin and hemin, also stabilizes the fast-inactivated state, which constitutes one of the prerequisites to obtain the effect of deltamethrin on Nav. Moreover, we report that under treatment with octopamine (1 µM) in the presence of PBO, deltamethrin can further decrease the sodium current amplitude (Extended Data fig. 6). This effect is not observed with a phosphatase inhibitor okadaic acid, reinforcing the role played by the phosphatase PP1/2A in the perturbation of the cytochrome P450 conformational landscape. This transient reshape of the active site conformation could explain an effective PBO displacement from the active-site cavity unmasking and orienting heme to facilitate the electrostatic interactions between the positively charged heme^72^ and the positively charged residues of Nav. This efficiently optimizes cytochrome P450-Nav interactions to stabilize Nav inactivated-state, which is essential for the action of deltamethrin. The immunocytochemical studies confirmed a tight coupling of activities between cytochrome P450 and Nav, where the spatial overlap between Nav and the cytochrome P450 CYP4 was revealed. By contrast, a low concentration of octopamine (100nM), which corresponds closely to the level of octopamine released under chemical stress condition *in vivo*^56^, acts through Gαs activated PKA-dependent pathway and phosphorylates DARPP-32, which blocks PP1-2A. The resulting inhibition of the cytochrome P450 dephosphorylation reduces the deltamethrin effect on Nav. The change in cytochrome P450 conformation, which may stabilize PBO in the active site cavity and masking the electrostatic repulsive interactions between the iron-containing porphyrin and the Nav positive charges exposed to the cytosol. This effect explain why PBO hampers the effect of deltamethrin on Nav.

Finally, we demonstrate the physiological relevance of these results. Octopamine, one of the main biogenic amines identified in insects, is released under stressful conditions ^51^. Previous reports indicate that stress conditions are closely associated with either an increase in the tyramine β-hydroxylase activity, which catalyzes the last step in octopamine biosynthesis or the enhancement of the octopamine levels^73,74^. We show that sublethal doses of the insecticide bendiocarb, used as a chemical stressor, elevated the octopamine level *in vivo*. This results in a decrease in the deltamethrin efficacy, as expected from our *in vitro* experiments performed with a physiological concentration of octopamine (i.e., 100nM).

In conclusion, we have described the complex molecular mechanism explaining why the deltamethrin effect on Nav is hampered when used with the common synergist, PBO. Octopamine, released under stress conditions, acting *via* the α-adrenergic-like octopamine receptors, triggers a biochemical pathway that modulates the cytochrome P450. A reduction of P450 activity (either by PP1/2A or binding of PBO), in turn, affects Nav channel gating and the final effect of pyrethroids. These could be explained by the disruption of the stabilization of the inactivated state of Nav, which is critical for deltamethrin to reach its final binding site in the DII-DIII fenestration and to exert the full insecticidal effect. Taken together, we reveal a novel adaptation mechanism to stressful conditions, such as exposure to sublethal doses of insecticides. The presented mechanism, likely to operate in other insect species, increases insect survival by reducing insecticide efficacy. This study reinforces the use of an alternative pyrethroid formulation technology, such as nanocapsules carrying deltamethrin, proving a higher efficiency *in vivo*^48,75^ than classical formulations containing pyrethroids combined with PBO. A deeper understanding of pesticides and synergists modes of action can support the development of more effective formulations and technologies for vector control, helping to optimize resources and maximize impact.

---

**Extended DataTable 1.** Electrophysiological characteristics of activation and inactivation of Nav.

|  | Activation |  | Inactivation |  |
| --- | --- | --- | --- | --- |
|  | V<sub>0.5</sub> (mV) | k (mV) | V<sub>0.5</sub> (mV) | k (mV) |
| **Control** | -21.6 ± 2.3 (n=6) | 5.2 ± 0.3 (n=6) | -34.1 ± 3.4 (n=5) | -8.5 ± 0.4 (n=5) |
| **PBO** | -32.9 ± 3.1 (n=7) | 4.6 ± 1.3 (n=7) | -35.7 ± 2.1 (n=7) | -5.7 ± 0.6 (n=7) |
| **DM** | -35.1 ± 3.5 (n=5) | 3.5 ± 0.7 (n=5) | -55.0 ± 2.0 (n=4) | -6.7 ± 0.4 (n=4) |
| **PBO + DM** | -38.5 ± 2.7 (n=6) | 5.2 ± 0.9 (n=6) | -45.4 ± 3.8 (n=5) | -6.3 ± 1.0 (n=5) |
| **Hemin** | -34.0 ± 2.5 (n=5) | 4.8 ± 1.0 (n=5) | -49.7 ± 2.4 (n=5) | -6.0 ± 0.1 (n=5) |
| **Oct (100nM)** | -36.4 ± 2.1 (n=6) | 2.4 ± 0.6 (n=6) | -41.7 ± 2.3 (n=5) | -6.2 ± 0.3 (n=5) |
| **Oct (1μM)** | -46.9 ± 1.8 (n=4) | 2.2 ± 0.5 (n=4) | -55.4 ± 2.4 (n=6) | -6.4 ± 0.2 (n=6) |
| **PBO + Oct (1μM) + DM** | -50.6 ± 3.5 (n=5) | 2.5 ± 0.5 (n=5) | -61.3 ± 2.3 (n=5) | -5.7 ± 0.5 (n=5) |
| **PBO + OkAc + Oct (1μM) + DM** | -39.4 ± 5.3 (n=4) | 5.0 ± 1.5 (n=4) | -55.1 ± 2.1 (n=8) | -5.9 ± 0.2 (n=8) |

**Extended Data Table 2.** Residues of the insect voltage-gated sodium channel PaNav1 found to contribute to deltamethrin binding. Residues are renumbered based on the house fly sodium channel (*Musca domestica*, GenBank accession number: X96668). For the references to the knockdown resistance (kdr) conferring mutations see the review (https://doi.org/10.3390/insects13080745) and additional publications (https://doi.org/10.1186/s13071-019-3565-x, https://doi.org/10.1371/journal.pntd.0004696).

| Channel segment | *Musca domestica* | *Periplaneta americana* | *kdr* |
| --- | --- | --- | --- |
| **DI S6** | V410 | V420 | + |
| **DII, S4-S5 linker** | M918 | M905 | + |
| **DII S5** | L932 | L919 | + |
|  | I936 | I923 | + |
| **DII P** | F979 | F966 | + |
|  | L982 | L969 | + |
|  | C983 | C970 |  |
| **DII S6** | N1013 | N1000 | + |
|  | V1016 | V1003 | + |
|  | L1017 | L1004 | + |
|  | F1020 | F1007 | + |
| **DIII P** | F1496 | F1497 |  |
| **DIII S6** | I1532 | I1533 | + |
|  | I1533 | I1534 | + |
|  | F1534 | F1535 | + |
|  | F1537 | F1538 | + |
|  | L1540 | L1541 |  |

**Supplementary Table 1.**
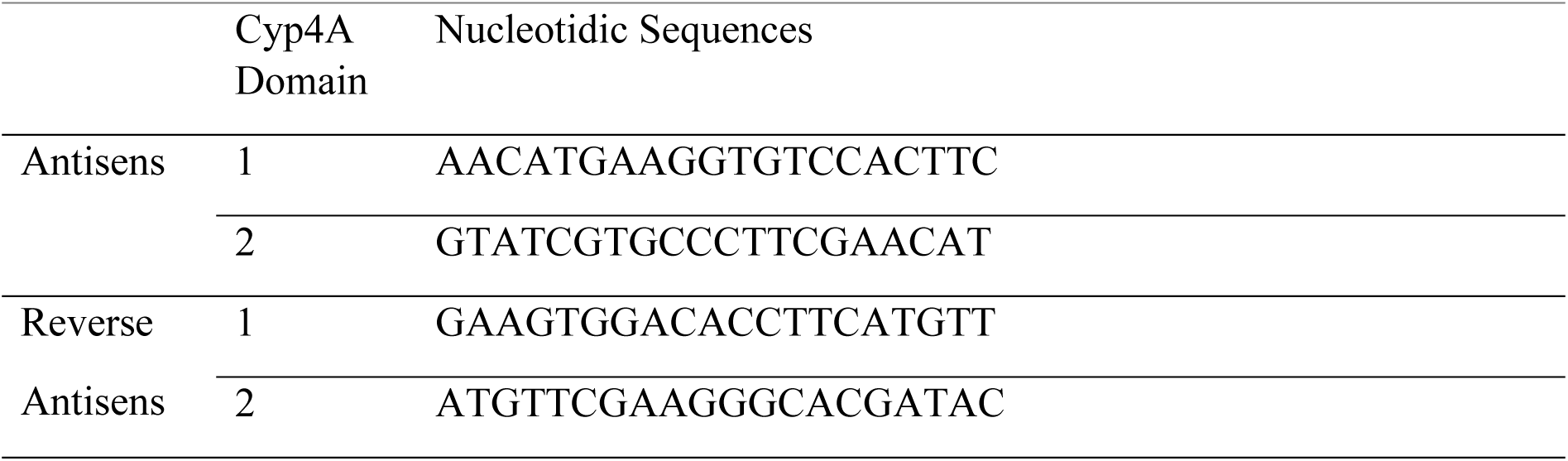
Nucleotidic sequences used to design antisens oligonucleotides and reverse antisens oligonucleotides.

## Methods

### Mosquito strains and bioassays

Bioassays were conducted using the standard WHO cylinder test procedure^33^ with few modifications to allow the assessment of the potentiation effect of the synergist PBO on pyrethroids against *Ae. aegypti* mosquitoes^76^. Briefly, the method consisted of exposing insects (i.e. 3-5 days old, unfed females for mosquitoes) for 1h to PBO at a concentration of 0.1–4% before exposing them for 1 h to Whatman no. 1 filter papers impregnated at the expected LC50 or below of a given pyrethroid (i.e. the type II alpha-cypermethin). Mortality of insects was recorded 24h post exposure to the insecticide. Testing and holding temperature and relative humidity were 27°C ± 2°C and 75 ± 10%, respectively. Whatman papers were impregnated according to the WHO standard procedure^33^. PBO was provided free of charge by Endura, India, whereas the insecticide alpha-cypermethrin was provided by BASF, Germany. Bioassays were conducted against a single strain of *Aedes aegypti* (Rockefeller) strains, reared at the Swiss Tropical and Public Health Institute (Swiss TPH), Basel, Switzerland and the Instituto Oswaldo Cruz (IOC) Fundação Oswaldo Cruz (FIOCRUZ), Rio de Janeiro, Brazil. The colonies were maintained in an insectarium for >15 generations and were fully susceptible to insecticides.

### DUM neuron isolation procedure

Experiments were carried out on Dorsal Unpaired Median (DUM) neuron somata isolated from the midline of the terminal abdominal ganglion (TAG) of the nerve cord of adult male cockroaches *Periplaneta americana* as already described^77^. Animals were immobilized ventral side up on a dissection dish. The ventral cuticle and the accessory gland were removed to allow access to the TAG, which was carefully dissected under a binocular microscope and placed in normal cockroach saline containing (in mM): 200 NaCl, 3.1 KCl, 5 CaCl_2_, 4 MgCl_2_, 10 HEPES, and 50 sucrose, pH was adjusted to 7.4 with NaOH. Isolation of adult DUM neuron somata was performed under sterile conditions by using enzymatic digestion by collagenase (type IA, 280 IU/ml; Worthington Biochemicals, Freehold, NJ) at 29°C for 35 min. A mechanical dissociation through fire-polished Pasteur pipettes was then used to isolate DUM neurons from the TAG. DUM neuron somata were maintained at 29°C for 24h before experiments were carried out.

### Calcium imaging

Isolated DUM neuron cell bodies were obtained, as already mentioned above. The cells were washed two times in saline and incubated in the dark with 5µM Fura-2 pentakis (acetoxy-methyl) ester (Fura-2 AM) (Sigma-Aldrich, Saint Quentin Fallavier, France) in the presence of 0.1% pluronic acid F68 (Sigma-Aldrich, Saint Quentin Fallavier, France) for 1h at 37°C. Pluronic acid is a nonionic surfactant used as a stabilizer of cell membrane protecting from membrane shearing to facilitate uptake of Fura-2 AM. After loading, cells were washed two times in saline. The glass coverslips were then mounted in a recording chamber (Warner Instruments, Hamden, CT, USA) connected to a gravity perfusion system allowing drug application. Imaging experiments were performed with an inverted Nikon Eclipse Ti microscope (Nikon, Tokyo, Japan) equipped with epifluorescence. Excitation light was provided by a 75-W integral xenon lamp. Excitation wavelengths (340nm and 380nm) were applied using a Lamdba DG4 wavelength switcher (Sutter instrument, Novato, CA, USA). Images were collected with an Orca-R2 CCD camera (Hamamatsu photonics, Shizuoka, Japan) and recorded on the computer with Imaging Workbench software (version 6, Indec BioSystems, Santa Clara, CA, USA). Experiments were carried out at room temperature in saline without sucrose. Intracellular calcium level was expressed as the ratio of emitted fluorescence (340/380 nm), as previously reported.

### Antisense Assay

Antisense oligonucleotides were designed to target the CYP4 family P450. Human CYP4A11 protein sequence (NCBI-Gene ID 1579) was blasted against *Periplaneta americana* transcriptome shotgun assemblies (GAWS, GEIF, GBJC) using tblastn. Top hit results were selected and corresponding nucleotide sequences were translated to identify the characteristic CYP4 family 13-amino acid motif, EVDTFMFEGHDTT. Nucleotide sequences were aligned and 20-mer antisense oligonucleotides (ASO) were designed to target the CYP4 consensus sequence in *Periplaneta americana*. To inhibit CYP4 expression, isolated DUM neurons, obtained as described just above, were incubated for 24h and 48h at 29°C with a 10 µM mixture of two 20-mers ASO (Eurogentec, Seraing, Belgium) targeting the 39 bases of CYP4 consensus sequence mRNA (Supplementary Table 1). Control experiments were performed using a mixture of reverse ASO (Supplementary Table 1).

### Electrophysiology- Whole-Cell Recordings and Data Analysis

A whole-cell patch-clamp recording configuration was used to record voltage-dependent sodium currents (voltage-clamp mode) and action potentials (current-clamp mode) at room temperature (20–22°C) in isolated DUM neuron cell bodies according to the protocol described just above. Patch pipettes were pulled from borosilicate glass capillary tubes (GC 150T-10; Clark Electromedical Instruments, Harvard Apparatus, Edenbridge, UK) using a P-97 Flaming/Brown Micropipette Puller (Sutter Instrument Company, Novato, U.S.A). Pipettes had resistances ranging from 0.8 to 1.2 MΩ when filled with internal pipette solution (see composition below). The liquid junction potential between bath and internal solutions was always corrected before the formation of a gigaohm seal (>2 GΩ). Voltage-dependent sodium currents and action potentials were recorded with an Axopatch 200A (Molecular Devices, Sunnyvale, CA) amplifier, filtered at 5 kHz using a 4-pole low-pass Bessel filter and were low-pass filtered at 10 kHz with clampfit software (version 10.1; Axon instruments).

For voltage-clamp experiments, DUM neuron cell bodies were voltage-clamped at a steady-state holding potential of −90 mV, and 10-ms test pulses were applied at a frequency of 0.2 Hz (except where otherwise stated). Although leak and capacitive currents were compensated electronically at the beginning of each experiment, subtraction of residual capacitive and leakage currents was performed with an online P/4 protocol provided by pClamp. The voltage-dependence of the conductance was calculated as a function of the membrane potential according to equation (1) and was fitted according to the Boltzmann equation (2):

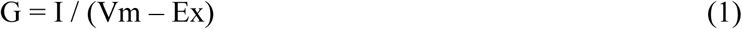

Where G is the sodium conductance, I is the inward current amplitude, Vm is the potential at which the membrane is clamped and Ex is the equilibrium potential for a given ion.

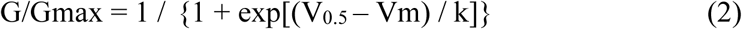

where G is the sodium conductance, Gmax is the maximal conductance from a holding potential of −90 mV, Vm is the membrane potential applied, V_0.5_ is the potential at which half the sodium channels are activated and k is the slope factor. Conductance-voltage relationships were fit for each individual cell. The voltage-dependence of steady-state inactivation was also examined using a double-pulse protocol by holding the cells at prepulse potentials between −100 and +40 mV, in 10-mV increments, for 500 ms. Thereafter the membrane potential was stepped back to the holding potential (−90 mV) for 1 ms before a 10-ms test potential to 0 mV. The inactivation curves were constructed by plotting the normalized amplitude of the peak current against the conditioning potential. Average data were best fitted according to the Boltzmann distribution (3):

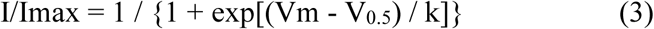

where I is the peak current, Imax is the maximal peak current from a holding potential of −90 mV, V_0.5_ is the potential at which half current was inactivated, Vm is the potential of the conditioning pulse and k is the slope factor. Inactivation curves were fit for each individual cell. We also examined the use-dependent action of deltamethrin. Use-dependent block was explored by 10 pulses applied at 10 Hz in DUM neuron held at −90 mV to 0 mV (pulse duration 2.5 ms). The peak current during each depolarization was normalized to the peak current during the first depolarization and plotted against the depolarization number.

### Solution and drug applications

Pharmacological agents were applied by a gravity perfusion valve controller system (VC–6 M, Harvard apparatus, 1 s in duration) controlled by pClamp software (flow rate of perfusion: 0.5 mL/min). The perfusion tube was placed within 100 μm from the isolated neuron cell body. For voltage-clamp experiments, the extracellular solution contained (in mM): 40 NaCl, 140 TEA-Cl, 3.1 KCl, 2 CaCl_2_, 7 MgCl_2_, 1 CdCl_2_, 5 4-aminopyridine (4-AP), and 10 HEPES buffer; pH was adjusted to 7.4 with TEA-OH. Patch pipettes were filled with internal solution containing (in mM): 90 CsCl, 70 CsF, 15 NaCl, 1 MgCl_2_, 5 EGTA, 3 ATP-Mg, and 10 HEPES buffer; pH was adjusted to 7.4 with CsOH. For current-clamp experiments, bath solution superfusing the cells contained (in mM): 200 NaCl, 3.1 KCl, 5 CaCl_2_, 4 MgCl_2_, and 10 HEPES; pH was adjusted to 7.4 with NaOH. Patch pipettes were filled with solution containing (in mM): 160 K/Dgluconate, 10 KF, 10 NaCl, 1 MgCl_2_, 0.5 CaCl_2_, 1 ATP; 0.1 cAMP, 10 EGTA, and 10 HEPES; pH was adjusted to 7.4 with KOH. Drug solutions were prepared in the external solution and applied in the immediate vicinity of the cell by a gravity perfusion system. In some cases, the tested compounds were added in the internal pipette solution immediately before use. In addition, some pharmacological agents were prepared in DMSO and then diluted in the bath solution to obtain the different concentrations tested. The highest concentration used in the electrophysiological recordings of DMSO was 0.1%. This concentration of solvent was not found to have any effect on the electrophysiological properties of DUM neuron cell body.

### Statistics and reproducibility

Data analysis and fitting procedures were performed with Prism v8 (GraphPad Software, Inc, San Diego, CA). Data are presented as the mean ± S.E.M. When data followed a normal distribution, significant differences were assessed with Student t-tests for multiple comparisons. In other cases, significant differences were assessed with Mann-Whitney tests. Statistical analysis was expressed as non-significant for *P* ≥ 0.05 and significant for \**P*< 0.05, \*\**P*< 0.01,

\*\*\**P*< 0.001 and \*\*\*\**P*<0.0001

### Immunocytochemistry

For light microscope immunocytochemistry, isolated DUM neurons were fixed for 50 minutes with 2 % paraformaldehyde containing 5 % (m/v) sucrose in phosphate-buffered saline (PBS, pH 7.4). After fixation, cells were washed three times for 5 min each in PBS and 5 min in PBS containing 0.2% Triton X-100 (PBS-T), respectively. To block non-specific binding of the primary antibodies, cell bodies were preincubated with 4% bovine serum albumin (BSA) in PBS-T for 1 h. Primary antisera rabbit polyclonal anti-voltage-gated sodium channel (SP19 Segment) antibody (Sigma Chemicals, L’isle d’Abeau Chesnes, France) or mouse monoclonal anti-Cyp4A11 (Bio-Techne Ltd., Abingdon, UK), both diluted 1/100 in PBS-T, were applied overnight at 4°C. After repeated washing in PBS-T, the secondary antibody, fluorescein isothiocyanate-labeled goat anti-rabbit IgG (FITC) or Alexa Fluor-labeled anti-mouse IgG (Thermo Fisher Scientific Inc., Illkirch-Graffenstaden, France), diluted 1/250, in PBS-T containing 4% BSA, were applied at 20°C for 3 h in the dark. Isolated DUM neuron cell bodies were then rinsed in 4% BSA in PBS and mounted on glass slides in glycerol/PBS. The control preparations were treated identically, except that the controls included the omission of primary antibodies. Preparations were viewed using a Nikon A1 confocal laser scanning microscope (x4 objective), Nikon A1 HD25. The images were acquired using 477 nm and 638 nm laser for excitation for FITC and Alexa Fluor, respectively. The emissions were collected *via* a photomultiplier through band-pass filter at 511 nm and 685 nm for FITC and Alexa Fluor, respectively. The images were processed using the NIS-Element (Nikon) and ImageJ (Fiji) software programs.

### Homology modelling

The homology models of an α subunit of the voltage-gated sodium channel protein PaNav1 from *Periplaneta americana* were built using the SWISS-MODEL server^78^, based on the D0E0C1_PERAM sequence from the UniProtKB. Models of closed, inactivated, and open structures were built using cockroach NavPas channel (PDB code: 6A95^79^), human hNav1.2 (PDB: 6J8E [10.1126/science.aaw2999]), and rat rNav1.5 (7FBS^80^) experimental structures, respectively. Two long intracellular loops (ICL1 – 330 residues, ICL2 – 260 residues) were removed from each model. The quality was validated using ERRAT^81^, PROCHECK^82^, and PROVE^83^, implemented in the Protein Structure Analysis and Verification Server provided by UCLA-DOE, Institute for Genomics and Proteomics. Further protein preparations including protonation state assignment, hydrogen addition, and minimization were conducted with Schrodinger Maestro^84^. The ERRAT overall quality factor, expressed as the percentage of the protein for which the calculated error value falls below the 95% rejection limit, equals 95.09 for the open model. Residues exceeding the error value were found in loop regions. The PROVE validation was passed with 0% buried outlier protein atoms. 93.4% of residues were found in the PROCHECK most favored regions. The electrostatic potential was calculated using APBS-PDB2PQR software.

### Molecular docking

3D structures of deltamethrin, PBO, and TTX were downloaded from the Cambridge Crystallographic Data Centre (accession number 1985206)^85^, and PubChem^86^, respectively. Molecular docking was performed using the smina package^87^. Ten independent runs (giving up to 100 poses each) of flexible ligands docking to the rigid protein with default parameters were conducted. For deltamethrin, additional 10 rounds of docking with the flexible sidechains option (--flexres) were performed, and the makeflex.py Python script provided with docking package was used to build the full receptor with the flexible residues re-inserted. Flexible sidechains were selected based on the best-scored positions of deltamethrin docking to the rigid protein using a 5Å cutoff.

### Classical molecular dynamics

Inputs for MD simulations were generated using CHARMM-GUI builder^88^. A heterogeneous bilayer model composed of 400 lipid molecules in proportions: 38% DOPE, 18% DOPS, 16% DOPC, 13% POPI, 11% SM (CER180), 3% DOPG, and 1% PALO 16:1 fatty acid were built to mimic an insect-like membrane. Water molecules were added above and below the lipids to generate a 20 Å thickness layer. The system was neutralized with Na^+^ and Cl^−^ ions to the concentration of 0.15 M. The CHARMM36 force field^89^ with the TIP3P model for water was applied. Six steps of equilibration composed of NVT dynamics (constant particle number, volume, and temperature) followed by NPT dynamics (constant particle number, pressure, and temperature) with gradually decreased restraint force constants to various components were run using NAMD 2.13^90^. Independent replicas of production MD simulation were run for each system: 4×500 ns for an open model, 3×500 ns for the open model with deltamethrin docked, and 3×500 ns for the inactivated model. The temperature was controlled by the Langevin thermostat with a value of 303.15 K. The target pressure was set to 1.01325 bar (1 atm). Topology and parameters files for deltamethrin were generated by SwissParam^91^.

### Enhanced sampling MD

Enhanced sampling simulations were run using the Gromacs 2020.7 software^92^ patched with the PLUMED 2.8 plugin^93^. Voltage sensors were removed to decrease the system size. The CHARMM36 force field and a time step of 2 fs were used. The simulations were run in the NVT ensemble using the stochastic velocity rescaling thermostat at 300 K with a relaxation time of 1 ps. We constrained hydrogen bonds using LINCS^94^. To find ligand dissociation pathways, well-tempered metadynamics- was used to bias the distance between the centers of geometries of the ligand and the protein; 15 trajectories were launched from the position of the ligand in the deep pose (p1) and 15 trajectories with the ligand in the shallow pose (p2). Gaussians were deposited every 1 ps along the distance using a height of 1 kJ/mol, a width of 0.02 nm, and a bias factor of 5. The dissociation process was terminated when the ligand reached the interior of the protein channel or the membrane. The minimal distance between any heavy atoms of the ligand and the protein was estimated by the radius of the tunnel using the maze plugin from PLUMED [https://doi.org/10.1016/j.cpc.2019.106865]. The free-energy profiles of the dissociation process were reconstructed by running two 300 ns well-tempered metadynamics simulations. The parameters were identical as for sampling the dissociation pathways. The simulations were started from the shallow binding pose (p2). For the dissociation pathway to the interior of the channel and the membrane, the distance was restrained to sample the ranges of 0.4 nm and 1.7 nm, and 1.6 nm and 3 nm, respectively. The free-energy profiles were reweighted using the Tiwary-Parrinello algorithm^95^, taking into account the metadynamics bias potential and the restraints.

### Fenestration analysis

MoleOnline with default parameters was used to visualize tunnels and to create a hydrophobicity-charge map of the static open PaNav1 model. The bottlenecks of channel fenestrations in static models, as well as the dynamical changes in their diameter based on the open model MD trajectories, were calculated using CAVER 3.0^96^. NAMD trajectories were processed into a series of PDB snapshots with a time step of 1 ns. For the analysis, the lipid bilayer, water molecules, ions, voltage-sensors, and ICLs were removed from the system so that calculations were performed on the pore domain helices (S5-S6 helices and P-loops), together with S4-S5 linkers and ECLs only. As starting points for tunnel search, given residues were selected from S6 helices of each domain: Phe417 (DI), Gly999 (DII), Phe1538 (DIII), and Phe1837 (DIV). Default parameters were used, except for a larger spherical probe radius and the shell depth of a surface region, both set to 15 Å.

### Cockroach motor ability tests, toxicity tests and spectrometry

Adult male cockroaches (*Periplaneta americana*) were obtained from Nicolaus Copernicus University breeding facility. The insects were kept at 29 ± 1 °C, fed with oat flakes, dog chaws, and water ad libitum.

Exposure to chemicals; bendiocarb (used in the form of insecticide Ficam 80 WP; Bayer, Poland) was dissolved in 96% ethanol to the concentration of 10 nM immediately before the experiment and a 100-fold dilution in water was made to obtain a final concentration of 0.1 nM. For the control group, a 100-fold dilution of 96% ethanol was made to obtain the same concentration as in bendiocarb solution (vehicle). The exposure to 0.1 nM bendiocarb was performed in a glass containers (12 cm diameter, 10 cm height) on 5 cockroaches by spraying them with 1 mL of solution. Control insects were sprayed with the vehicle. Insects were exposed for 45 min for octopamine level evaluation and 60 min for toxicity tests. Deltamethrin was dissolved in DMSO to 10 mM concentration immediately before the experiment. Then, 100-fold dilutions with water were made to obtain final concentration of 1 µM. The bottom of a glass container was lined with paper, and 1 mL of deltamethrin solution was spread directly before experiment. Five insects were moved from 0.1 nM bendiocarb exposure to the container with the desire deltamethrin concentration.

Motor ability test; to observe the paralysis caused by deltamethrin exposure, a test for motor abilities was performed. After pre-exposure with 0.1 nM bendiocarb and 1-hour exposure with 1 µM deltamethrin, insects were placed on the dorsal side on the cork arena (50 cm diameter) and recorded with video-camera (Logitech C920) at 30 f/s speed. Each cockroach was placed in exactly the same way by only one experimenter. The time of turning back from the dorsal to ventral side was evaluated by counting the time on the video where the insect was freely released and the time at which insects reached a 90° turnover. For toxicity test, the mortality of insects was assessed. After pre-exposure with 0.1 nM bendiocarb, insects were exposed to deltamethrin for 48 hours. Insects were recorded with video-camera (Logitech C920), and every 30 minutes, a 5-second video was captured. The time of death was evaluated, and insects were considered dead when they remained on dorsal side without any movement. Kaplan-Meier survival analysis was then performed.

Liquid chromatography tandem-mass spectrometry LC-MS/MS; Insects were placed in individual 100 mL containers and left to adjust for 5h. Then, without handling the insects were sprayed with 0.1 nM bendiocarb or the vehicle. After 45 minutes, cockroaches were sacrificed by microwave irradiation (800 W, 30 s). Individual cockroaches were weighted, fragmented and placed in the 5 mL Eppendorf tube with 2 mL of water. Using an ultrasound homogenizer (VCX-130), samples were carefully homogenized, and 666 µL of concentrated HCl was added for protein precipitation. Homogenates were centrifuged (10 min, 5,000 g; Eppendorf 5804-R) and 650 µL of supernatant was carefully collected and placed in fresh 2 mL tubes. Samples were centrifuged again (10 min, 14,000 g; Eppendorf 5804-R), and 350 µL of the supernatant was collected for examination. To adjust pH of samples, 150 µL of 8.5 M NaOH was added together with 5 µL of d_3_-octopamine solution (as internal standard, 1 µg/mL). The level of octopamine was analyzed using LCMS-8045 tandem mass spectrometry (Shimadzu Corp.) based on preliminary analytical conditions designated with DL-octopamine hydrochloride standard (Sigma-Aldrich, Poland) and DL-octopamine-d_3_ hydrochloride (ChemCruz, Santa Cruz Biotechnology Inc.). Accucore™ Amide HILIC, 2.6 µm, 2.1 mm × 100 mm HPLC column equipped with a precolumn was used to perform chromatographic separation. Two solutions were used as the mobile phase: A) 25 mM ammonium formate with 0.05% formic acid and b) 85% acetonitrile with 0.05% formic acid (v/v). The parameters of separation were as follows: a linear gradient of 90–50% of B for 4.5 min, flow rate of 0.4 mL/min, 35 °C. Negative electrospray ionization (ESI) was used in mass spectrometry, ions were fragmented by CID- collision-induced dissociation at 3 kV. The presence of octopamine was confirmed with multiple reactions monitoring (MRM transitions 136.1–92.3 and 136.1–65.2 for octopamine and 139.1–93.3 and 139.1–67.3 m/z for d_3_-octopamine).

### Statistical analysis

For the comparison of data with non-normal distribution, the Kruskal–Wallis test was used, followed by Mann–Whitney post-hoc test. For only two sample comparison, the t-test or Mann-Whitney test were performed. Normality of data was tested with Shapiro-Wilk test. The analysis was conducted in the IBM SPSS 25 Statistics software (IBM Corporation, Armonk, NY, USA). The results were expressed as mean values ± SE. The differences were considered significant when \**P* < 0.05.

## Data availability

The data that support the computational part of the study are available to download from GitHub at https://github.com/jakryd/nav-deltamethrin-md

The data that support this study are available from the corresponding authors upon request.

## Acknowledgements

We thank Fabienne Simoneau, Aurélia Rolland and Prof. David Macherel for their skillful assistance at the IMAC-SFR QUASAV, Angers, France for light microscope immunocytochemistry facility. We thank Julie-Anne Hugel, Salome Keller and Julian Adler for insects rearing, Martin Gschwind, Xavier Nicolas, Théo Homet and Capucine Legendre for their technical assistance. This research was funded by the National Science Centre, Poland, under grant no. 021/41/N/NZ3/02165 (B.N.). Computations were carried out using the computers of Centre of Informatics Tricity Academic Supercomputer & Network (CI TASK).

## Author contributions

B.L. and W.N. conceived, designed, and directed the study; B.N. created homology models of the cockroach PaNav1 sodium channel and performed molecular docking, classical molecular dynamics, and fenestrations analysis; B.L. and W.N. analyzed modeling results; J.R. performed enhanced sampling simulations; E.M. recorded and analyzed sodium channel currents in isolated cockroach DUM neurons and performed light microscope immunocytochemistry with the assistance of C.D and B.L.; C.D. designed antisense and reverse antisense oligonucleotides and performed calcium imaging experiments; V.C. and A.M. conducted bioassays on mosquito strains with the assistance of P.M.; M.J. conceived and conducted cockroach motor ability tests, toxicity tests, and spectrometry with the assistance of J.K.; B.L and B.N. wrote the manuscript with inputs from all authors.

## Extended Data Figure legends

**Extended Data Fig. 1.**
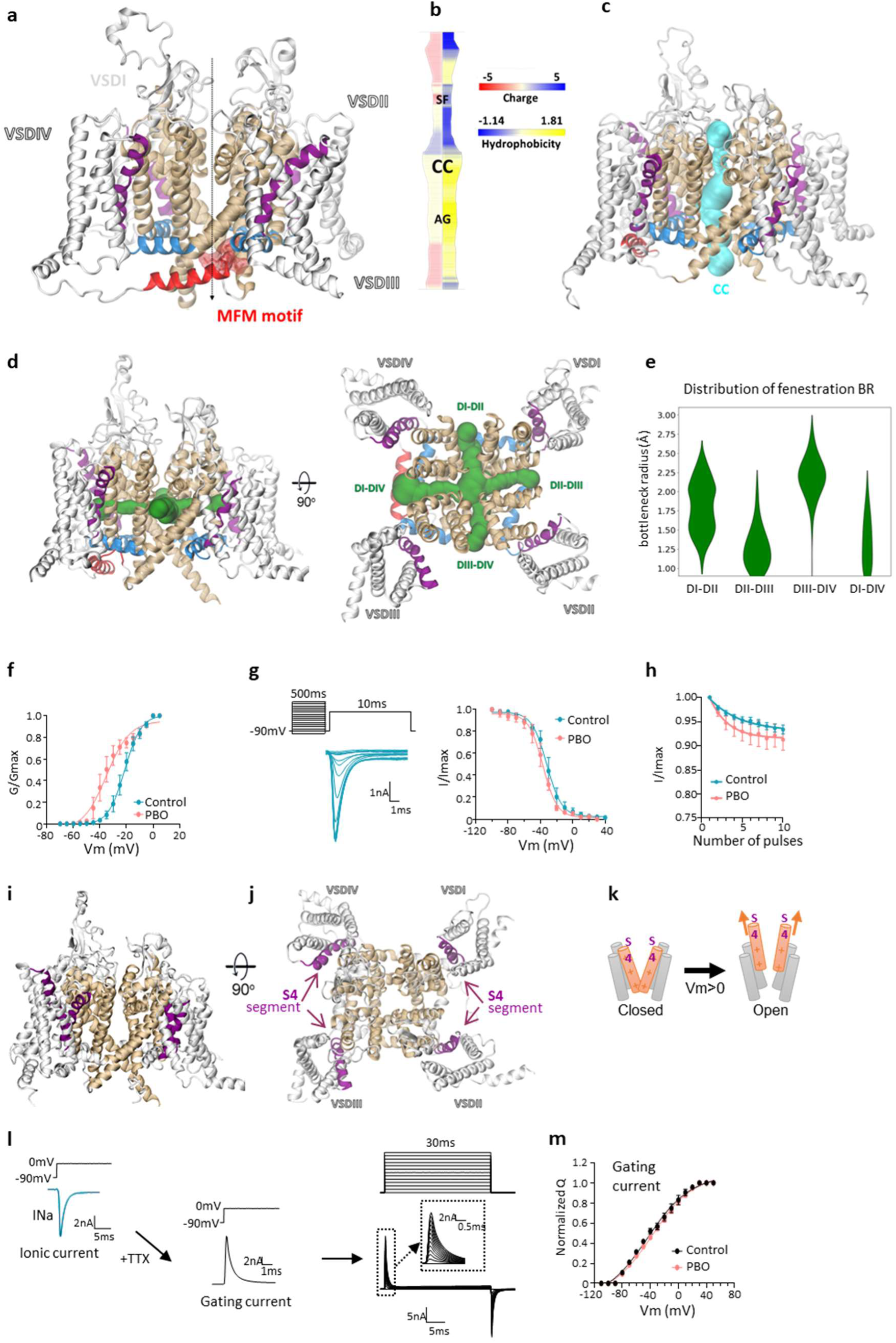
Voltage-gated Na channel structure and electrophysiological effects of PBO on the channel. **a.** Homology model of the cockroach PaNav1 sodium channel built based on a template of the open-state structure of rNav1.5 channel (PDB code: 7FBS). The α-subunit consists of a single polypeptide chain that folds into four domains (DI-DIV) built from six transmembrane helices (S1-S6) each. Helices 1-4 contributing to the voltage-sensor domain (VSD) with positively charged S4 helix (purple) act as voltage sensors, and helices S5-S6 create the pore domain (PD, gold). The S4-S5 linkers are in blue. The inactivation gate MFM motif, as a part of the DIII-DIV linker, is marked in red. An approximate location of the pore is indicated by the dashed line. **b.** The profile of the ion-conducting pore with the selectivity filter (SF), central cavity (CC), and activation gate (AG) regions marked. The left panel is colored according to the charge scale and the right panel represents the normalized hydrophobicity of the tunnel, ranging from the most hydrophilic (Glu=-1.140) to the most hydrophobic (Ile=1.810). The pore is visualized in cyan in (**c**). **d.** The side (left) and top (right) view of the channel with the four lateral fenestrations shown in green. **e.** The bottleneck radius distribution of the fenestrations based on three replicas of 500 ns classical molecular dynamics trajectories. **f.** G/Gmax *versus* potential of the sodium current recorded in control (n=6) and in the presence of 10 µM PBO (n=7), fitted with the Boltzmann equation (2, see Methods). **g.** Steady-state inactivation curves for sodium channels recorded in control (n=3) and with 10 µM PBO (n=9), averaged and plotted as a function of the prepulse potential. Smooth lines are fits to Boltzmann functions (3, see Methods). *Inset* shows typical sodium current recordings obtained according to the voltage protocol, as indicated. **h.** Use-dependant decrease in the sodium peak current amplitude in control (n=8) and with 10 µM PBO (n=7). **f-h.**; Data are means ± S.E.M.; **i-m.** Electrophysiological studies of the effect of PBO on sodium channel gating currents but not sodium current. **i-k.** Structure of the sodium channels, as explained in (**a**) representing the position of the positively charged S4 segments (purple) in both side (**i**) and top (**j**) views of the sodium channel. **k.** Scheme of the voltage sensing. Orange cylinders represent the positively charged S4 segment transitions in the channel during depolarization (Vm>0), producing the gating current. **l,m**. Gating currents were elicited by using a voltage–clamp protocol in the presence of 50 nM TTX (left panel). Depolarizing pulses were applied for 30 ms from −110 mV to + 50mV (middle panel), in control (n=5) and with 10 µM PBO (n=5). **m.** Q/V curve of peak gating current amplitudes. Data are means ± S.E.M..

**Extended Data Fig. 2.**
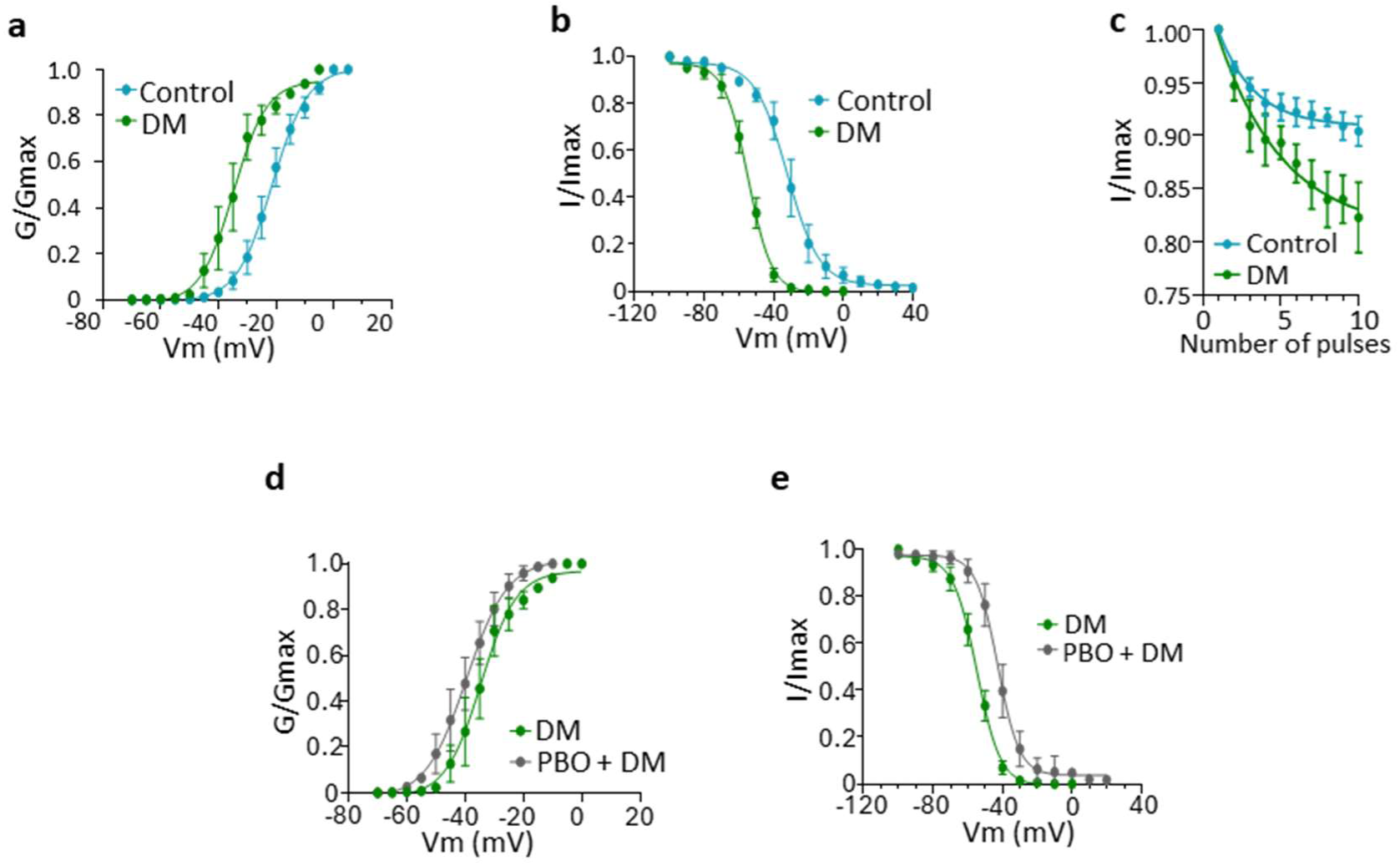
Biophysical action of deltamethrin on voltage-gated sodium channels. **a.** Normalized conductance–voltage (*G*–*V*) relations fitted with the Boltzmann equation (2) obtained in control (n=6) and in the presence of 1 µM DM (n=6). **b.** Voltage– steady-state inactivation relations (I/V) obtained from a three-pulse protocol established in control (n=3) and with 1 µM DM (n=4). Smooth lines are fitted to Boltzmann functions (equation 3). **c.** Use-dependant decrease in the sodium peak current amplitude in control (n=7) and with 1 µM DM (n=3). **a-c.** Data are means ± S.E.M.. **d**,**e.**. Normalized conductance–voltage (*G*–*V*) relations fitted with the Boltzmann equation (2) obtained in DM (1 µM, n=6) and with DUM neuron pre-treated with intracellular application of 10 µM PBO and then after in the presence of 1 µM DM (n=7) (**d**). Similar experimental conditions were used to study the voltage-dependence of the steady-state inactivation (**e**). Smooth lines are fitted to Boltzmann functions (3). Data are means ± S.E.M.

**Extended Data Fig. 3.**
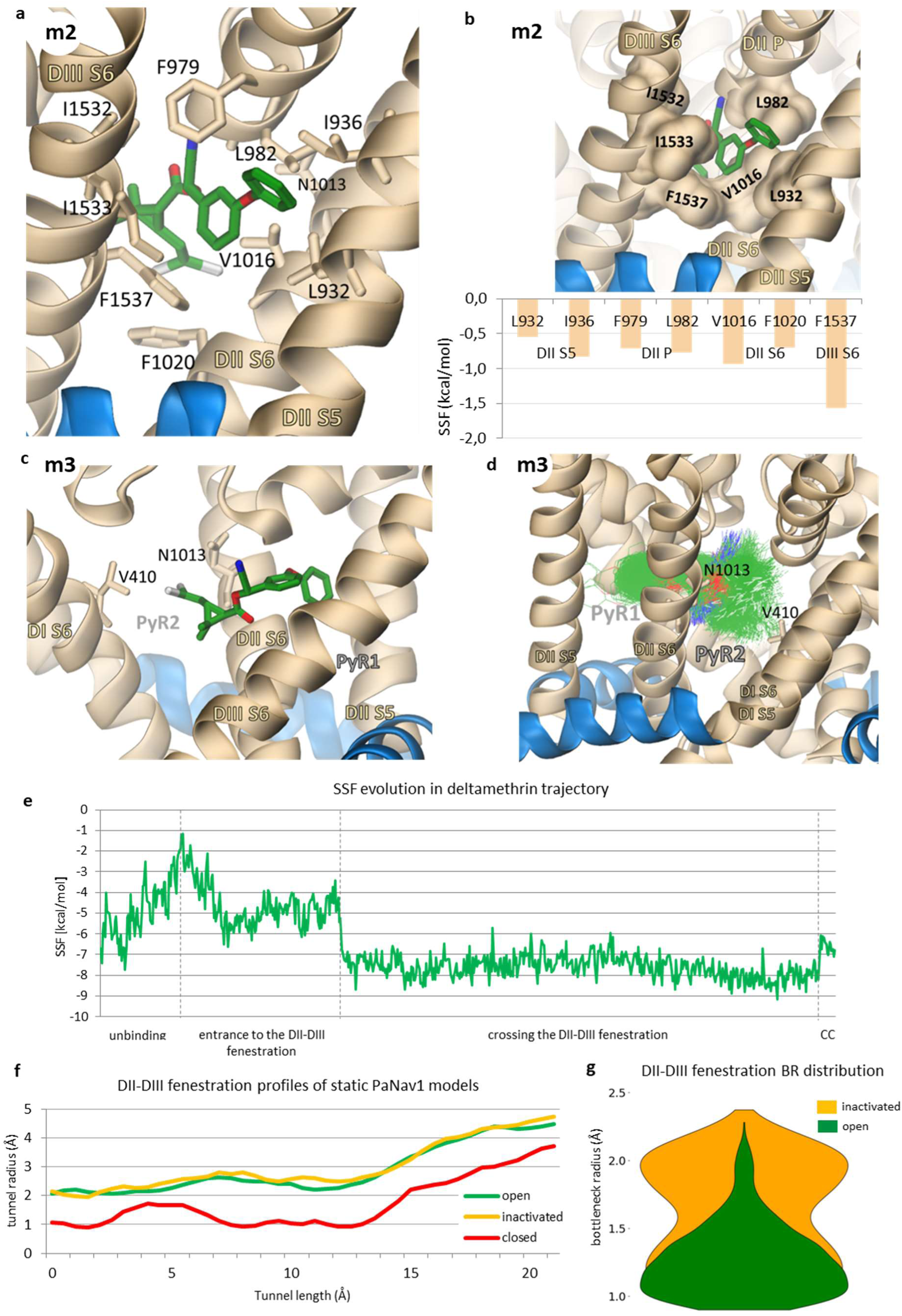
Deltamethrin interactions with insect voltage-gated sodium channel occur *via* fenestration – a molecular dynamics (MD) perspective. **a,b.** The detailed interaction analysis of deltamethrin in the best-scored pose from the MD simulations, corresponding to the free energy minimum 2. All marked residues are known to cause knockdown resistance when mutated (Extended Data Table 2). The fenestration-bottlenecking residues are presented in the surface representation in (**b**). The residues with the highest contribution to the binding energy, measured using smina scoring function (SSF), are shown in the bottom panel. **c,d.** Deltamethrin in a pose corresponding to the third free energy minimum (m3) interacts with the kdr residues, contributing to the previously proposed pyrethroids receptor site 2 (PyR2) while still maintaining contact with kdr residues from PyR1. An overlap of deltamethrin positions from the 3×500 ns MD is presented in (**d**). **e.** The deltamethrin-PaNav1 channel affinity is an example trajectory of ligand dissociation. The docking energy function SSF was used to calculate the interaction energy of the ligand moving through the fenestration. **f.** Comparison of the DII-DIII fenestration profiles in the homology models of PaNav1 channel – open in green, inactivated in orange, and closed in red, built based on experimental structures with the PDB code 7FBS, 6J8E, and 6A95, respectively. **g.** The bottleneck radius (BR) distribution of the DII-DIII fenestration. Data from 3×500 ns MD simulations were combined for each of the inactivated (orange) and open (dark green) models. Residues are renumbered based on the house fly sodium channel (*Musca domestica*, GenBank accession number: X96668).

**Extended Data Fig. 4.**
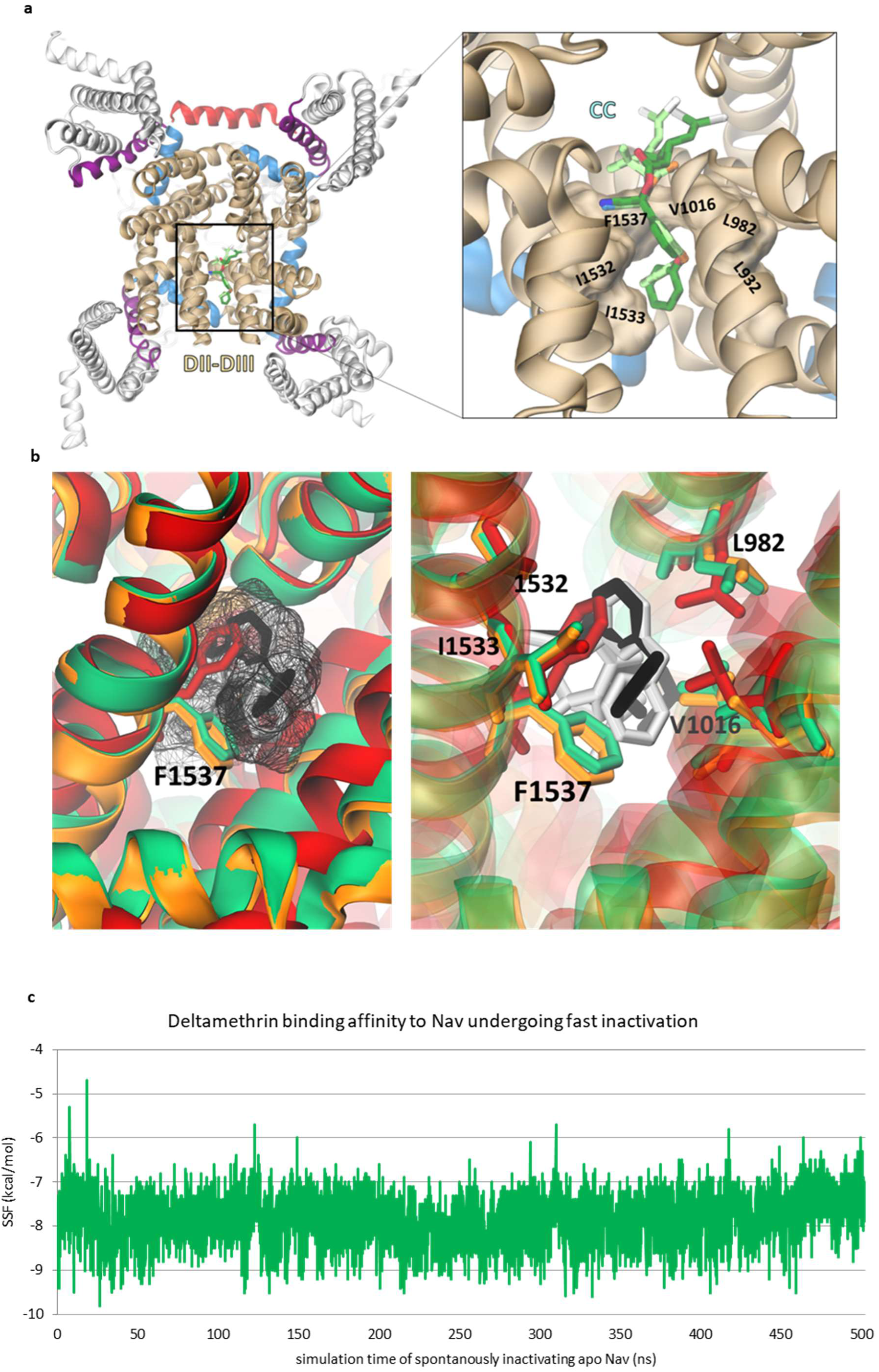
State dependence of deltamethrin binding. **a.** The lowest energy poses of deltamethrin docked to the open and inactivated state models of PaNav1 are shown in dark green and lime, respectively. The models were aligned, and the representation of the open model is shown for clarity (top view). **b.** Alignment of the open (green), inactivated (orange), and closed (red) state models (side view). The surfaces of deltamethrin bound to the open and inactivated models are shown in the left panel. Stick representation corresponds to the binding position in the open state (black) and inactivated state (silver). The residues creating free energy barriers in the DII-DIII fenestrations are marked. **c.** The binding affinity of deltamethrin and the inactivating channel calculated using smina scoring function (SSF) based on docking to the frames extracted from the 500 ns-long MD simulation.

**Extended Data Fig. 5.**
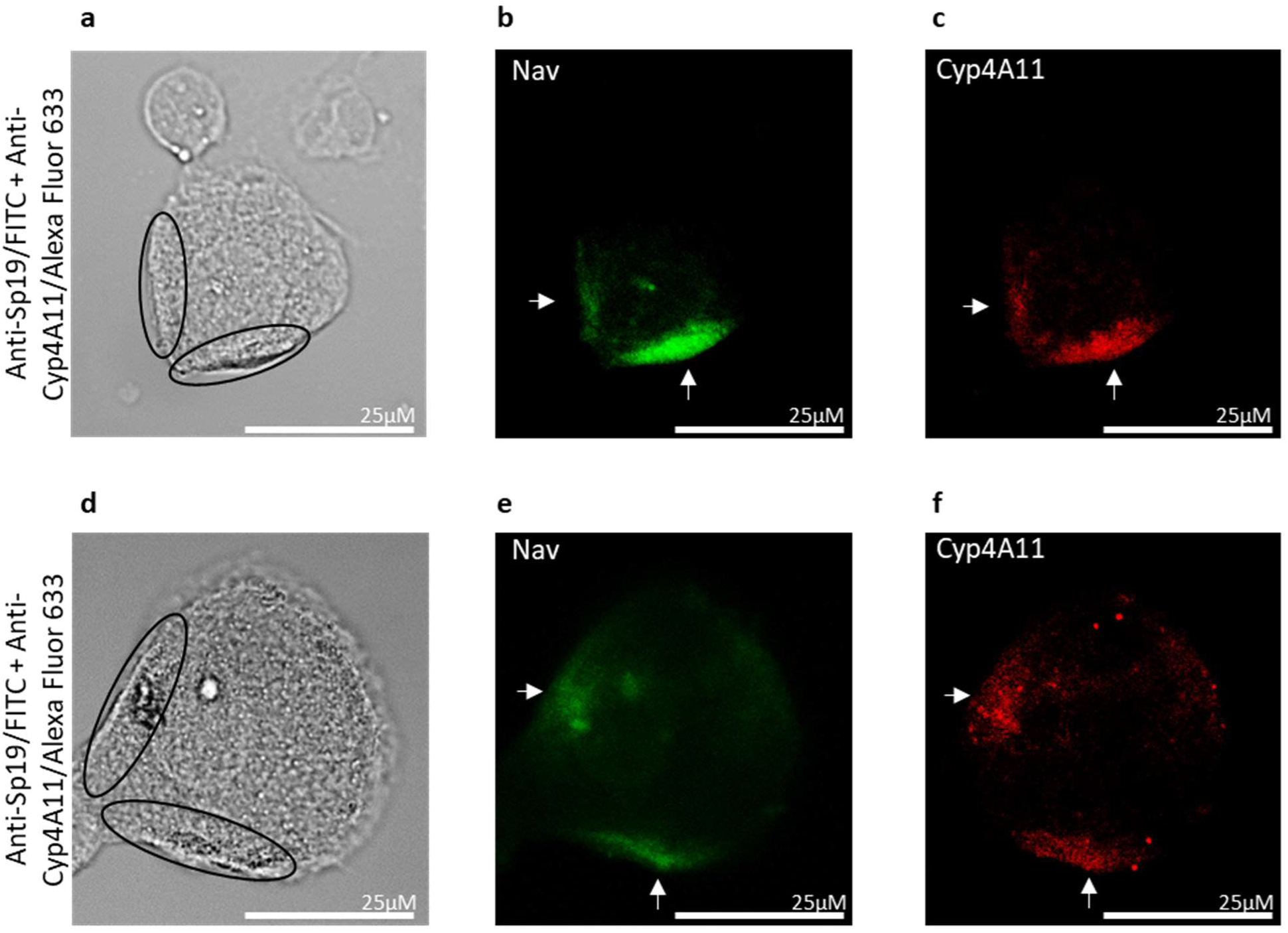
Immunocytochemistry showing co-localization of sodium channels and Cyp4A11 in isolated DUM neurons. **a,d.** Light photomicrographs of isolated adult DUM neurons. **b-f.** Isolated DUM neuron cell bodies labeled with anti-SP19 (Nav) antibody, which was detected with FITC conjugated secondary antibody (positive green fluorescence, white arrows) (**b,e**) and anti-Cyp4A11 antibody detected with Alexa Fluor 633 conjugated secondary antibody (positive red fluorescence, white arrows) (c,f). Both positive staining and co-localization of the two proteins present a granular appearance and is most intense in the same basal region of the soma close to the initial segment (open circles). Negative control experiments showing the specificity of the primary antibodies binding to the sodium channels and the Cyp4A11, together with the secondary antibody controls that show that the label is specific to the primary antibodies are illustrated in Supplementary Information Fig. 4.

**Extended Data Fig. 6.**
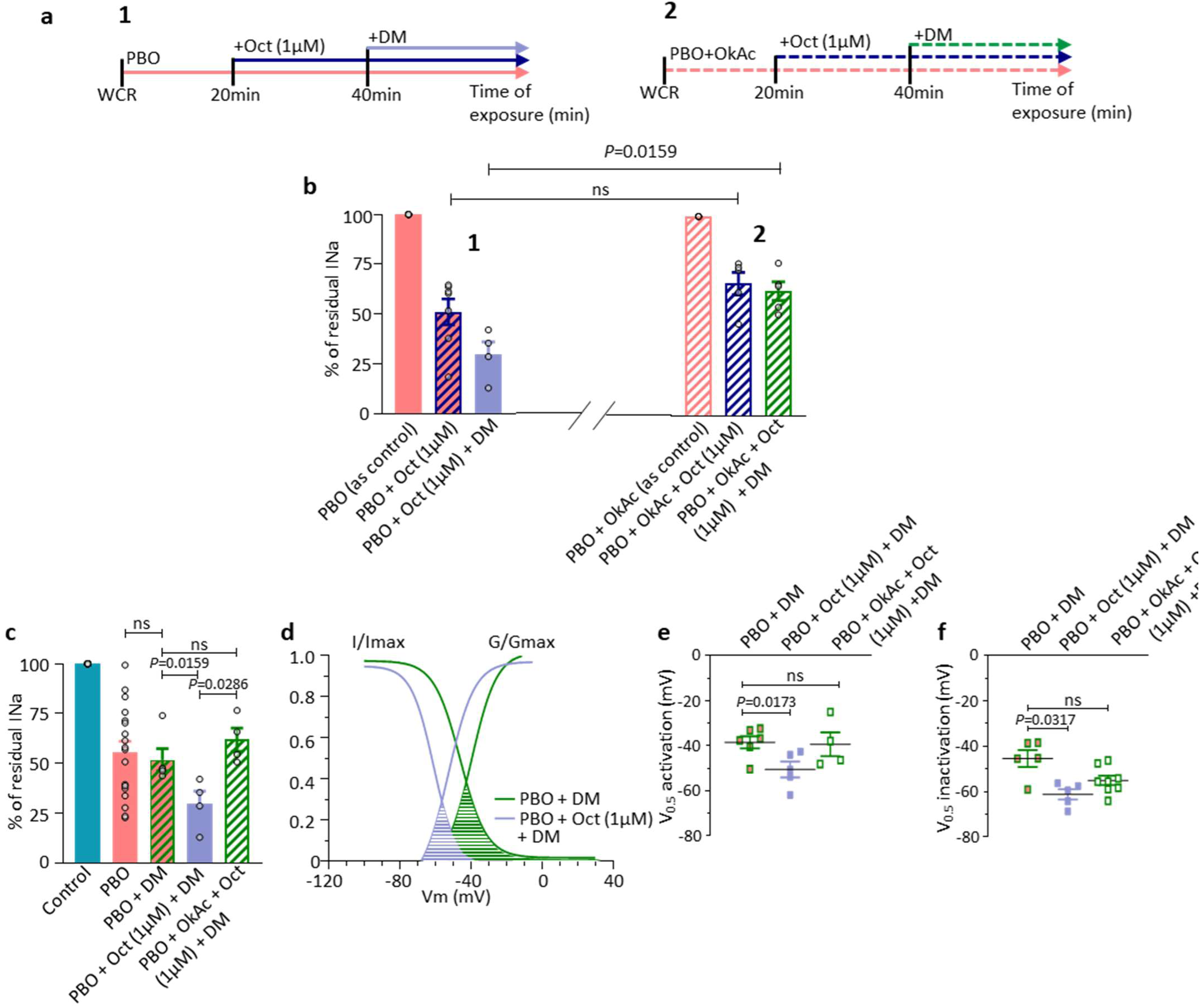
The dephosphorylation-induced cytochrome P450 conformational change affects the effects of PBO and deltamethrin. **a,b.** Comparative histogram illustrating the % of residual sodium current obtained at 0 mV from a holding potential of −90 mV recorded according to the protocols presented above (1 left and 2 right panels) (**a**). The sodium current is measured under different experimental conditions indicated below each bar (**b**). In the left panel (1), PBO (10 µM) is used as control (n=7), PBO (10 µM) + Oct (octopamine, 1 µM, n=7) and PBO (10 µM) + Oct (1 µM) mixed with DM (1 µM, n=4). In the right panel (2), PBO (10 µM) + OkAc (okadaic acid, PP1-2A inhibitor) (1 µM) is used as control (n=5), PBO (10 µM) + OkAC (1 µM) + Oct (1 µM, n=5), PBO (10 µM) + OkAc (1 µM) + Oct (1 µM) mixed with DM (1 µM) (n=5). **c.** Comparative histogram representing the % of residual sodium current recorded as indicated in (a), in different experimental conditions indicated below each bar, PBO (10 µM) (n=14), DM (1 µM) in the presence of PBO (10 µM) (n=5), PBO (10 µM) + Oct (1 µM) mixed with DM (1 µM) (n=4) and PBO (10 µM) + OkAc (1 µM) mixed with Oct (1 µM) and DM (1 µM) (n=4). **d.** Boltzmann functions describing the voltage dependences of opening (G/Gmax) and inactivation (I/Imax)) of sodium channels established with PBO (10 µM) + DM (1 µM) (n=6, green) and with PBO (10 µM) + Oct (1 µM) mixed with DM (1 µM) (n=5, blue). Colored areas indicate overlap between activation and inactivation curves, which is expected to result in the window current, which is larger for PBO + DM than PBO + Oct + DM. **e,f.** Scatter-plots showing the values of half-maximal voltage-dependence (V_0.5_) of activation (**e**) and inactivation (**f**) for sodium current in PBO (10 µM) + DM (1 µM) (n=6 and n=5, respectively), with PBO (10 µM) + (Oct, 1 µM) mixed with DM (1 µM) (n=5 and n=5, respectively) and with PBO (10 µM) + OkAc (1 µM) mixed with Oct (1 µM) and DM (10 µM) (n=4 and n=8, respectively). Data are means ± S.E.M.. P values are calculated by the Mann– Whitney U test.

**Extended Data Fig. 7.**
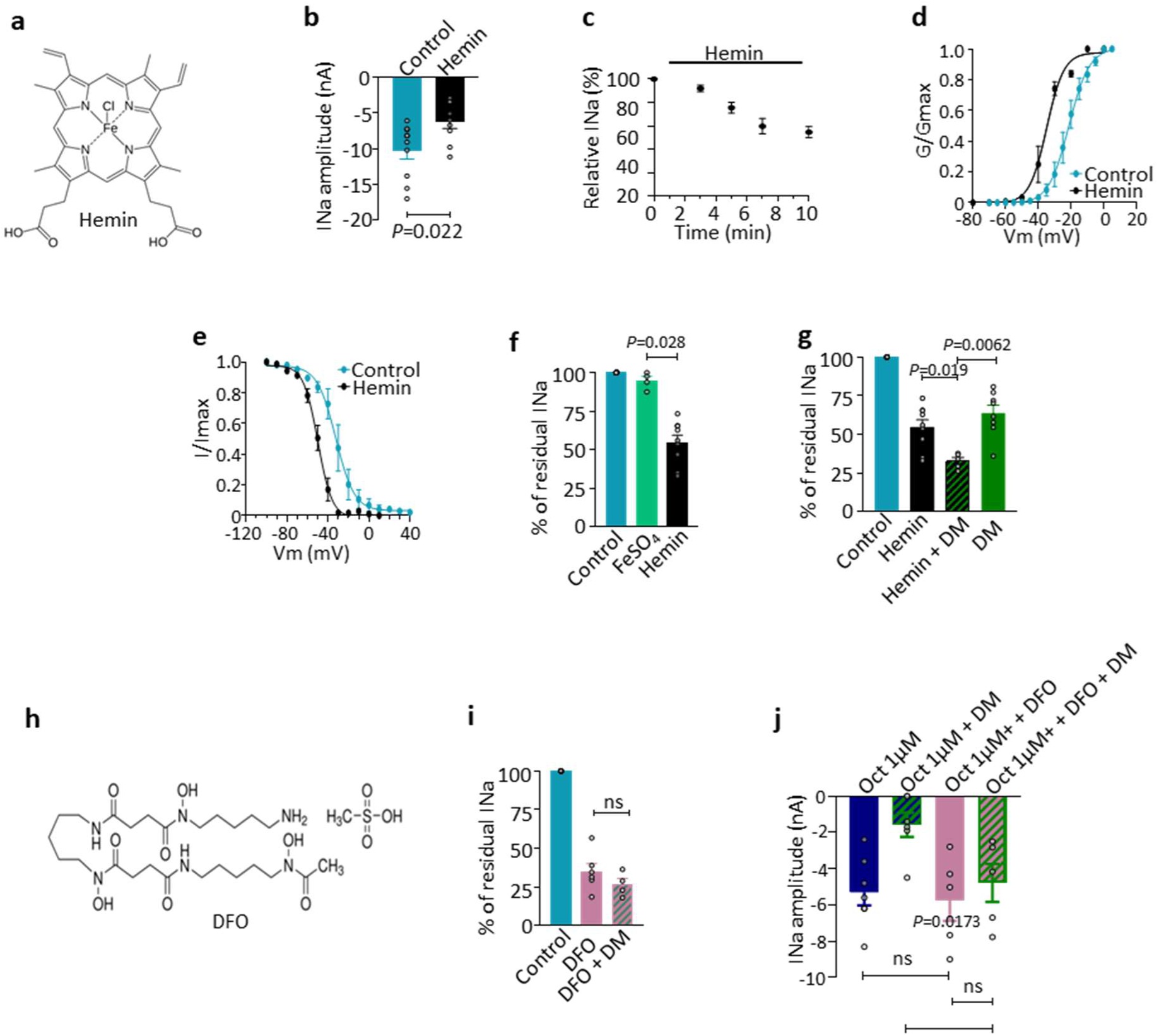
Intracellular hemin is a functionally effective endogenous modulator of sodium channel altering the effect of deltamethrin. **a.** Chemical structure of hemin. **b**. Comparative histogram of the sodium current amplitudes measured in control (n=10) and after intracellular application of 100 nM hemin (n=9). **c.** Relative peak sodium current (%) plotted *versus* different times, following the dialysis with 100 nM hemin. Time zero was marked at the time of rupture of the membrane patch for whole-cell recordings (n=4-10). **d.** G/Gmax *versus* potential of the sodium current recorded in control (n=6) and in the presence of 100 nM hemin (n=5), fitted with the Boltzmann equation (2). **e.** Steady-state inactivation curves for sodium channels recorded in control (n=5) and with 100 nM hemin (n=5), averaged and plotted as a function of the prepulse potential (see protocol in Extended Data Fig. 2g). Smooth lines are fits to Boltzmann functions (3). **f.** Comparative histogram showing the % of residual sodium current amplitude recorded in control (n=4), with 1 µM FeSO4 (n=4) and with 100 nM hemin (n=9). **g.** Residual sodium current (%) recorded in control (n=9), with 100 nM hemin (n=5) and with 1 µM DM in the presence of 100 nM hemin (n=5), compared to 1 µM DM applied alone (n=8). **h-j.** Comparative histogram illustrating the residual sodium current (%) measured in control (n=8), with DFO (deferoxamine mesylate, chemical structure in (**h**)), the iron chelator, wich interacts with hemin *via* the iron moiety) (10 µM) (n=7) and with DM (1 µM) in the presence DFO (10 µM) (n=4) (**i**). In all experiments, DFO was intracellularly applied. **j.** Comparative histogram illustrating the sodium current amplitudes recorded with Oct (1 µM, n=7), with Oct (1 µM) mixed with DM (1 µM, n=6), Oct (1 µM) in the presence of DFO (10 µM, n=5) and with Oct (1 µM) mixed with 1 µM DM, in the presence of 10 µM DFO, intracellularly applied (n=5). Data are means ± S.E.M.. *P* values are calculated by the Mann– Whitney U test.

**Extended Data Fig. 8.**
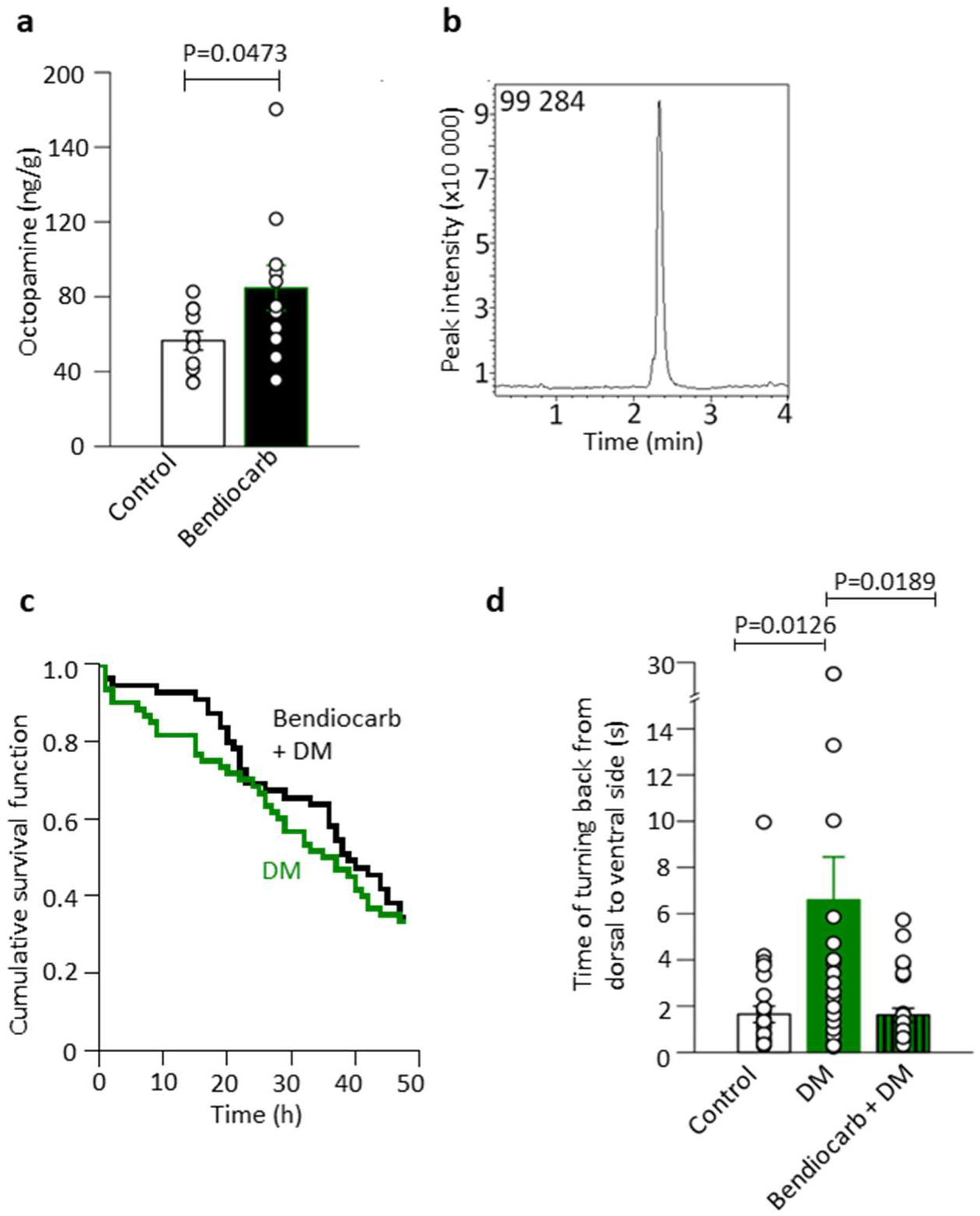
Chemical stress increases octopamine level in cockroaches *Periplanta americana* and decreases the effectiveness of deltamethrin. **a.** Octopamine level in the control group (white bar; n=11) and in insect exposed to 0.1 nM bendiocarb for 45 minutes (black bar, n=11). Data are means ± S.E.M.. *P* values are calculated by the Mann– Whitney U test. The difference was considered significant when \**P* < 0.05. **b.** Chromatogram for octopamine standard solution 1 ng/µl. The level of octopamine was analyzed using LC-MS/MS with octopamine-d_3_ as the internal standard. **c.** Cumulative survival analysis (Kaplan-Meier) for insects treated with 10 µM deltamethrin (green line, n = 60) and insects pre-exposed to 0.1 nM bendiocarb and then treated with 10 µM deltamethrin (black line, n = 55). **d.** Time of turning back from dorsal to ventral side of cockroach (motor ability test) for control (white bar, n=28), insects treated with 1 µM deltamethrin (green bar, n=26), and insects first pre-exposed to 0.1 nM bendiocarb and then treated with 1 µM deltamethrin (green striped bar, n=25). Data are means ± S.E.M.. *P* values are calculated by the Kruskal-Wallis test and Mann-Whitney U as a post-hoc test.

## Supplementary Figure legends

**Supplementary Fig. 1.**
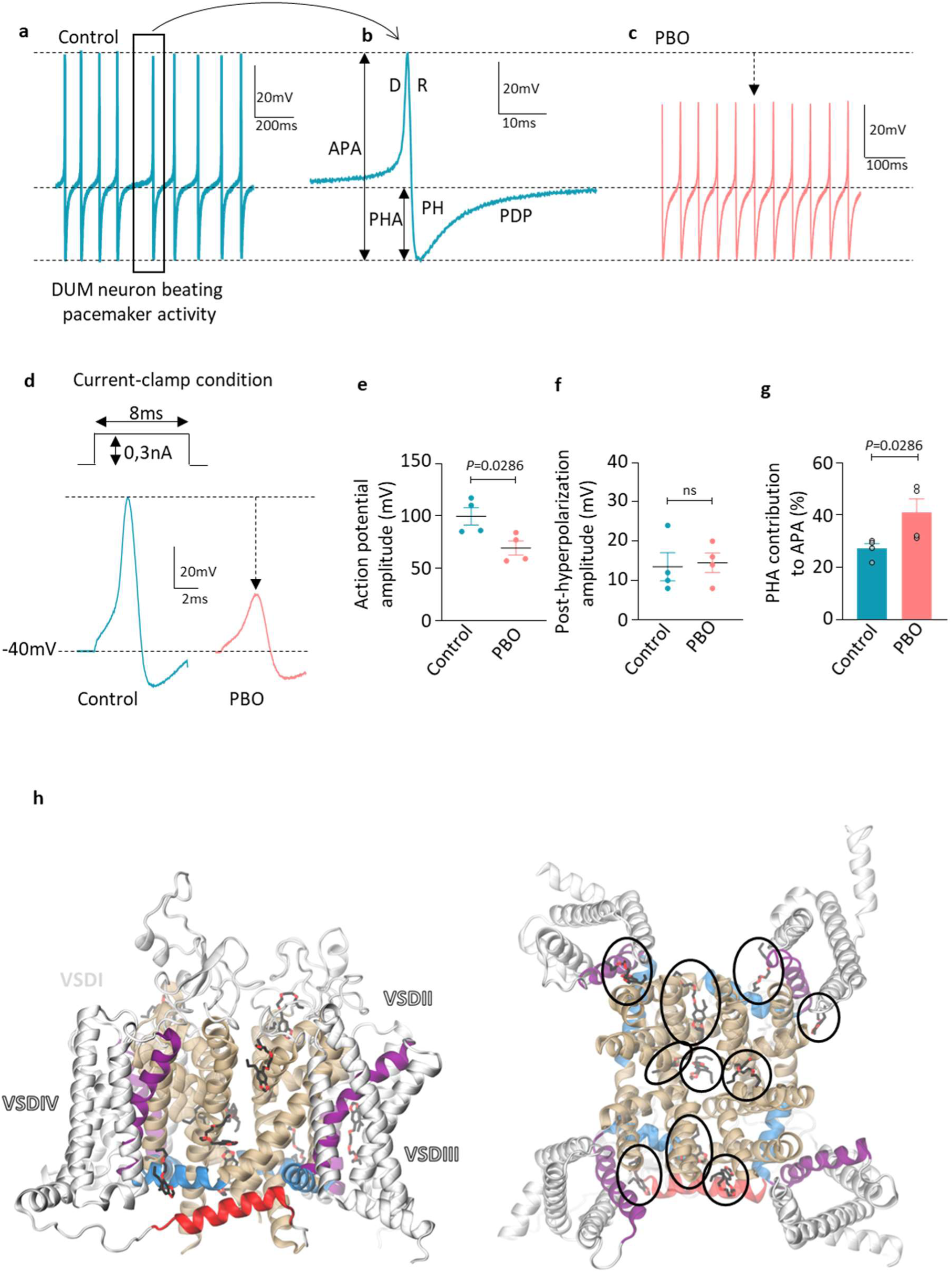
The cytochrome P450 inhibitor PBO alters the neuronal pacemaker activity. **a.** Example of intrinsic spontaneous overshooting action potentials expressed by an isolated DUM neuron cell body, recorded with the patch-clamp technique in the whole cell recording configuration under current-clamp condition. **b.** Representative action potential with the different phases of pacemaker activity including depolarization (D), repolarization (R), post-hyperpolarization (PH) and predepolarization (PDP). **c.** Representative neuronal pacemaker activity recorded in the presence of 10 µM PBO inducing specific reduction of the depolarizing phase due to an influx of sodium ions through voltage-gated sodium channels. **d.** Action potentials elicited under current-clamp condition according to the protocol indicated above traces, in control and in the presence of 10 µM PBO, intracellularly applied. **e,f.** Scatter plots showing the values of action potential amplitude (APA, **e**) and post-hyperpolarization amplitude (PHA, **f**), as indicated in (**b**) for action potentials recorded in control (n=4 and n=4, respectively) and in the presence of 10 µM PBO (n=4 and n=4, respectively). **g.** Comparative histogram representing the % of the PHA contribution to the APA measured in control (n=4) and 10 µM PBO (n=4). *P* values are calculated by the Mann–Whitney test, ns, non-significant (p>0.05). Data are means ± S.E.M.. **h.** The lowest-energy poses of PBO (shown in a black licorice representation) from 10 independent docking runs to the PaNav1 model in the open conformational state.

**Supplementary Fig. 2.**
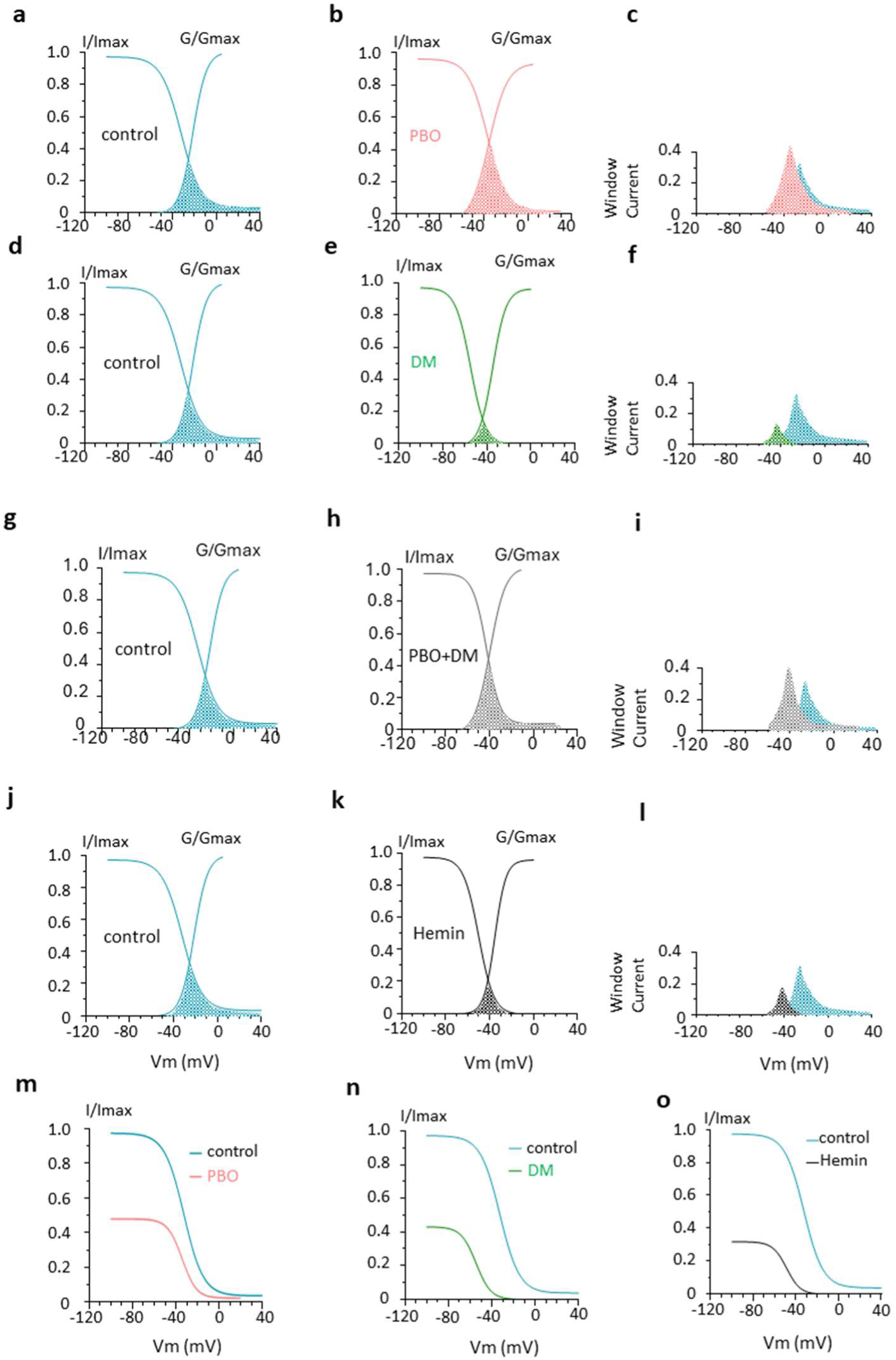
Window currents are affected by deltamethrin (DM), PBO and hemin. **a,d,g,j.** Continuous lines representing activation (G/Gmax) and steady-state inactivation (I/Imax), are fitted according to Equations (2,3) established in control (left column in blue). **b,e,h,k.** Middle column illustrating fits of activation (G/Gmax) and steady-state inactivation (I/Imax) established with PBO (10 µM, pink) (**b**), deltamethrin (DM, 1 µM, green) (**e**), PBO-DM (grey) (**h**) and hemin (100 nM, black) (**k**). **c,f,i,l.** Comparative superimposed estimated window currents (right column) corresponding to the overlaps between activation and inactivation curves under control and PBO (**c**), control and deltamethrin (**f**), control and PBO-DM (**i**) and control and hemin (**l**). **m-o.** Superimposed voltage-dependent inactivation curves representing plots of average normalized peak currents to their maximal value of current amplitudes *versus* prepulse potentials in control (blue) and in the presence of PBO (**m**), deltamethrin (DM, **n**) and hemin (**o**). Smooth lines are fitted to the Boltzmann function (3).

**Supplementary Fig. 3.**
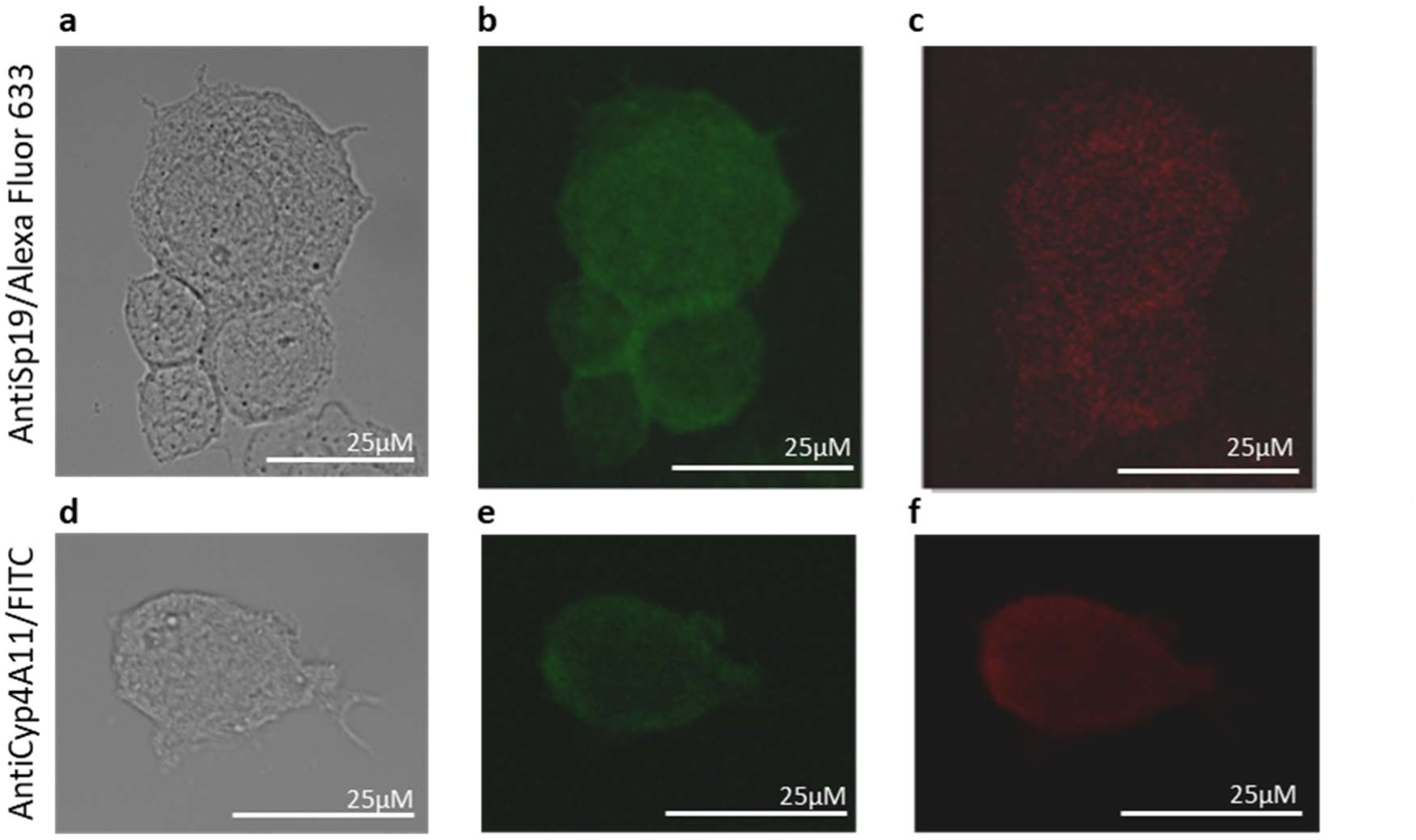
Negative controls for immunofluorescence. **a-c**. Light photomicrograph of isolated DUM neuron cell bodies (a), treated with anti-SP19 (Nav) antibody were incubated with Alexa Fluor 633 conjugated secondary antibody and recorded with the wavelength of excitation of FITC (477 nm), (**b**) and the wavelength of excitation of Alexa Fluor 633 (633 nm), (**c**). **d-f.** Isolated DUM neuron cell body (light photomicrograph (**d**), treated with anti-Cyp4A11 antibody were incubated with FITC conjugated secondary antibody and recorded with the wavelength of excitation of FITC (477 nm, **e**) and the wavelength of excitation of Alexa Fluor 633 (638 nm, **f**).

**Supplementary Fig. 4.**
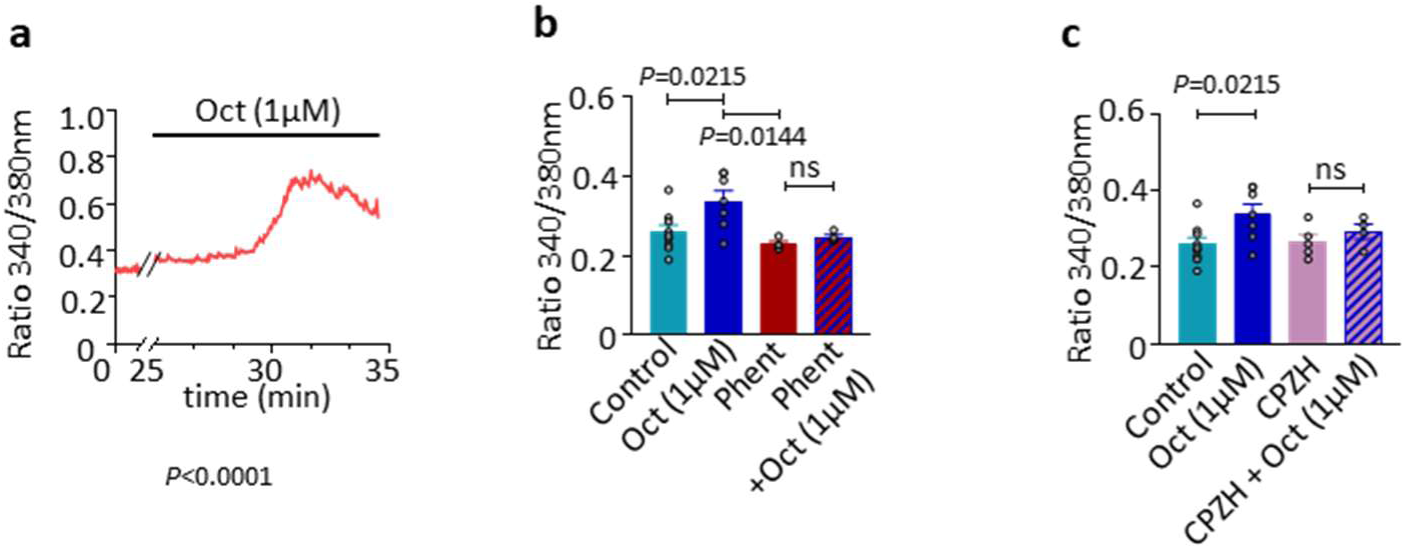
Octopamine exerts its effects through binding to G protein-coupled α-like adrenergic receptors (OctR α-AdrR). **a.** Bath application of 1 µM octopamine (Oct) increased intracellular free calcium concentration ([Ca^2+^]_i_). **b.** Comparative histogram illustrating the Fura-2 fluorescence ratio (340/380) of [Ca^2+^]_i_ recorded in control (n=9), with 1 µM octopamine (Oct, n=7), with phentolamine (10 µM Phent) alone (n=4) and phent (10 µM) mixed with Oct (1 µM, n=4). **c.** The effects of 1 µM Oct (n=7) was compared to the fluorescence ratio (340/380) recorded in control (n=9), with 500 nM CPZH (chlorpromazine, OctR α-AdrR antagonist) alone (n=5) and in the presence of 1 µM Oct (n=4). *P* values are calculated by the Student’s t test, ns, non-significant (p≥0.05). Data are means ± S.E.M..

**Supplementary Fig. 5.**
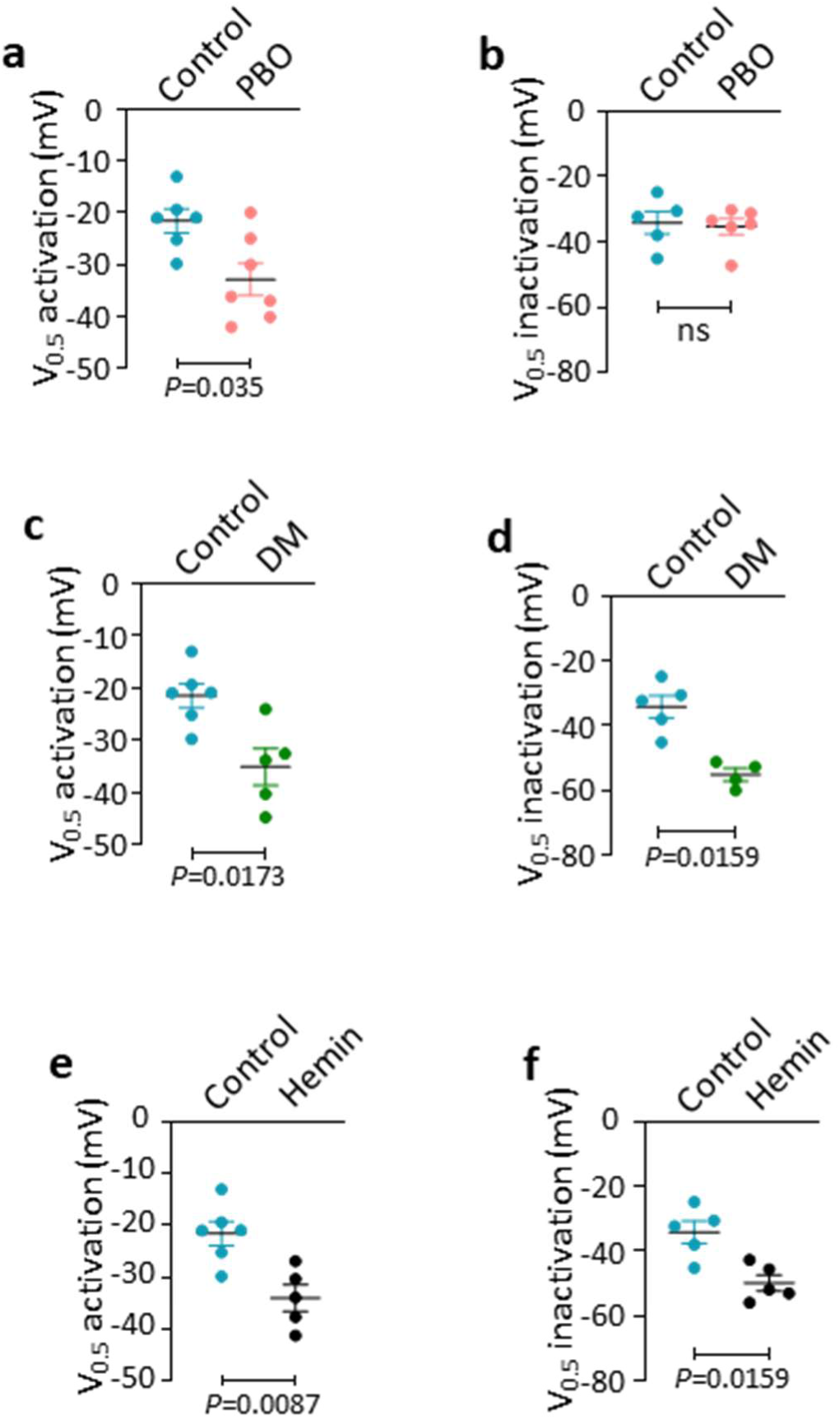
Scatter plots of half-maximal voltage-dependence of activation and inactivation. **a-f.** Scatter plots showing the values of half-maximal voltage-dependence (V_0.5_) of activation and inactivation for sodium current in control (n=6 and n=5, respectively) (**a**) and with 10 µM PBO (n=7 and n=6, respectively) (**b**). **c,d,** Typical scatter plots for half-maximal V_0.5_ activation and inactivation in control (**c**) (n=6 and n=5, respectively) and in the presence of 1 µM deltamethrin (DM) (n=5 and n=4, respectively) (**d**). **e,f.** Scatter plots showing the V_0.5_ activation and inactivation determined in control (n=6 and n=5, respectively) (**e**) and with 100 nM hemin (n=5 and n=5, respectively). *P* values are calculated by the Mann–Whitney test, ns, non-significant (p≥0.05). Data are means ± S.E.M..

**Supplementary Fig. 6.**
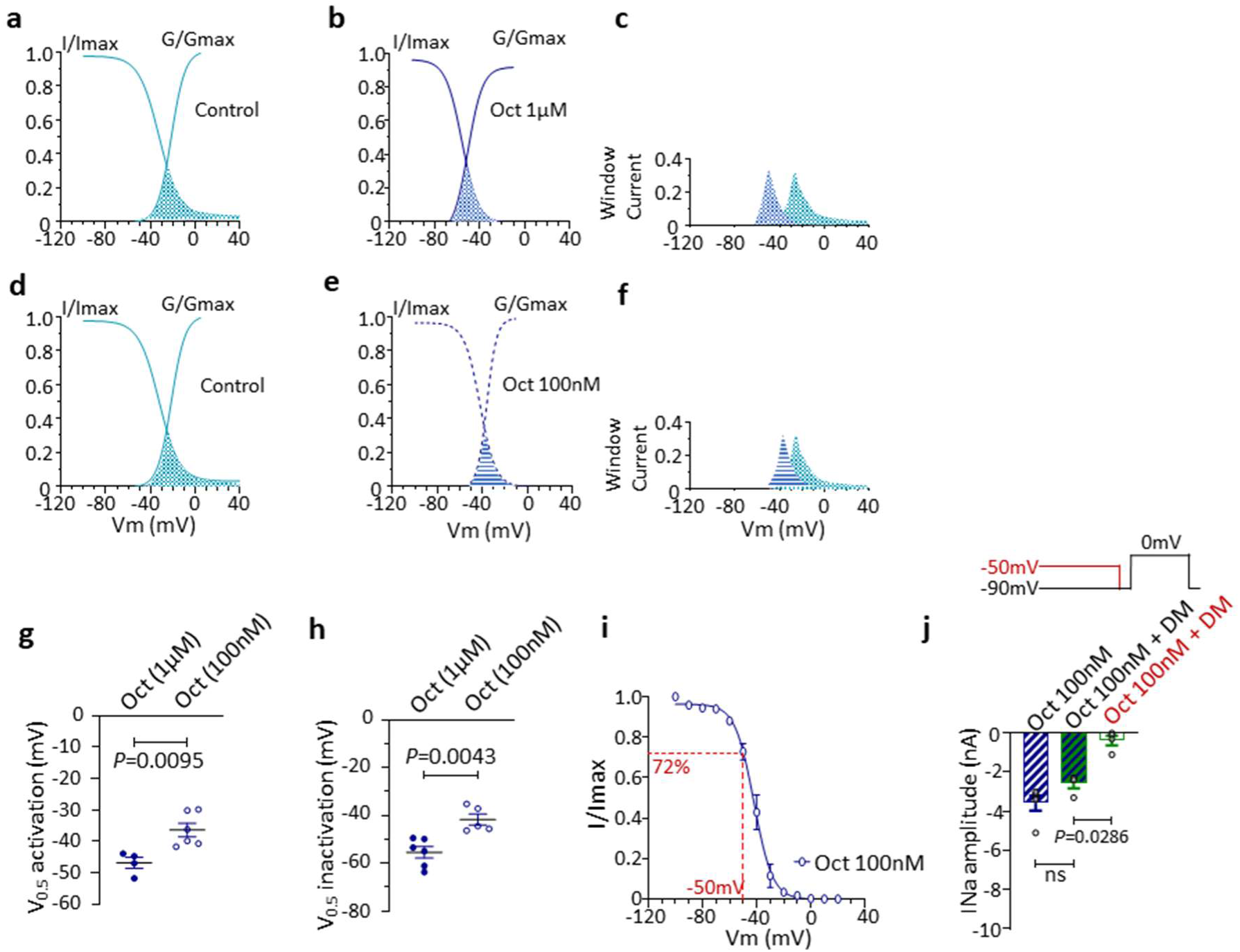
Concentration dependence of the effects of octopamine on voltage-gated sodium channel. **a-c.** Continuous lines representing activation (G/Gmax) and steady-state inactivation (I/Imax) are fitted according to Equations (2,3) established in control (n=6 and n=3, respectively) (**a**) and in the presence of Oct (octopamine) used at 1 µM (n=6 and n=6, respectively) (**b**). **c.** Superimposed estimated window currents corresponding to the overlaps between activation and inactivation curves. **d-f.** Fits of activation (G/Gmax) and steady-state inactivation (I/Imax) in control (n=6 and n=3, respectively) (**d**) and with lower concentration od Oct (100 nM) (n=6 and n=5, respectively) (**e**). **f.** Corresponding estimated window currents obtained as explained just above. **g,h.** Comparative scatter plots for half-maximal V_0.5_ activation between Oct (1 µM) (n=6) and Oct (100 nM) (n=6) (**g**) and inactivation between Oct (1 µM) (n=6) and Oct (100 nM) (n=5) (**h**). **i.** Steady-state inactivation curve for sodium channels recorded with 100 nM Oct (n=5), averaged and plotted as a function of the prepulse potential, according to the protocol shown in (**j**). Smooth line is fit to Boltzmann functions (3). **j.** Histogram representing the sodium current amplitudes recorded under different experimental conditions as indicated in *inset* and below each bar. Oct (100 nM, n=5), Oct (100 nM mixed with DM (1 µM, n=4) and Oct (100 nM mixed with DM, 1 µM) (n=4). In the last case, the sodium current was elicited at a holding potential of −50 mV, as indicated in red. *P* values are calculated by the Mann–Whitney U test. Data are means ± S.E.M..

